# eIF5A coordinates the transcription and translation of its target genes

**DOI:** 10.1101/2025.11.25.690413

**Authors:** Marina Barba-Aliaga, Lianqi Chi, Samoa Prieto-Díez, Jordi Planells, José García-Martínez, José E. Pérez-Ortín, Paula Alepuz

## Abstract

Maintaining balanced cellular protein levels requires precise control of gene expression and effective coordination between the various stages of the process, from transcription to translation. In recent years, several components of the translation apparatus have been found in the nuclei of various eukaryotes, where they regulate transcription, mRNA processing or export, thereby integrating different stages of gene expression. eIF5A is an essential and evolutionarily conserved translation elongation factor that is involved in viral infection and in the development of diseases such as cancer and neurodevelopmental disorders. eIF5A promotes translation elongation by binding to ribosomes that stall at codons encoding problematic amino acids for peptide bond formation, such as consecutive prolines, also known as polyproline motifs. Although eIF5A shuttles between the nucleus and cytoplasm, its specific nuclear roles remain poorly defined. Here, we demonstrate that nuclear yeast eIF5A binds to chromatin and represses gene transcription by preventing the binding of RNA polymerase II. Importantly, chromatin binding and transcriptional repression by eIF5A have a higher impact on genes encoding its own translational targets. The presence of polyproline motifs in genes imposes both translation and transcriptional control by eIF5A. Furthermore, eIF5A’s active engagement in cytoplasmic translation is necessary for its role in repressing transcription. Our results suggest that eIF5A coordinates gene expression by promoting the cytoplasmic translation of specific genes while repressing their transcription in the nucleus, thus ensuring efficient final protein synthesis.

**Significance Statement:** Our study provides genome-wide and gene-specific evidence supporting the role of the translation elongation factor eIF5A in transcription. eIF5A is essential in eukaryotes, facilitating the translation of mRNAs encoding stretches of problematic amino acids, such as consecutive prolines. Through its role in the synthesis of specific proteins, eIF5A has been linked to development and different diseases, including cancer and diabetes. We have now discovered that eIF5A also controls the transcription of its translation target genes and this effect is driven by the presence of eIF5A-dependent motifs at their sequences. In the nucleus, eIF5A binds to specific genes and attenuates the binding of RNA polymerase II. By negatively regulating transcription and positively regulating translation, eIF5A coordinates gene expression, fine-tuning protein levels.

## Introduction

Eukaryotic translation initiation factor 5A (eIF5A) is an essential, evolutionarily conserved protein with puzzling biological functions. eIF5A is the only known protein post-translationally modified by hypusination, which involves the addition of the polyamine spermidine via the action of two enzymes: deoxyhypusine synthase (DHPS) and deoxyhypusine hydroxylase (DOHH). eIF5A promotes cell proliferation and development, and is involved in apoptosis, autophagy and cytoskeleton organization, as well as maintaining mitochondrial activity. Non-physiological levels of eIF5A or its hypusination enzymes have been linked to diseases such as diabetes, chronic inflammation, altered immune responses, and various cancers. Furthermore, eIF5A promotes viral infection, and a decline in eIF5A levels has been linked to cellular ageing. Moreover, variants in eIF5A, DHPS and DOHH genes cause rare inherited neurodevelopmental disorders (see (1–7) for reviews).

Some of these diverse eIF5A biological functions rely on its role as a translation elongation factor, which is required for synthesizing specific proteins. Thus, although not necessary for each round of translation elongation, hypusinated-eIF5A binds to ribosomes and promotes the formation of peptide bonds between amino acids known to be poor substrates for this reaction that otherwise stall translation, such as consecutive prolines (polyPro motifs) but also combinations of proline, glycine and charged amino acids (8–10).

Despite being a translation elongation factor, eIF5A is not exclusively confined to the cytoplasm, as it shuttles between this compartment and the nucleus. In mammalian cells, the nuclear export of eIF5A preferably requires exportin 4 (Xpo4) (11, 12), and less frequently the nuclear export receptor Xpo1/CRM1 (13); whereas in yeast cells, the export relies on Pdr6 (14, 15). Remarkably, post-translational modifications dictate the subcellular distribution of eIF5A: hypusination and sumoylation promote cytoplasmic localization, while acetylation favors nuclear localization (16–18).

The partial nuclear localization of eIF5A is similar to that of other translation factors, which are among the most abundant proteins in the cell, and is likely to be related to additional functions in mRNA metabolism. In this regard, eIF5A is involved in the nuclear export of unspliced HIV-1 viral mRNA through its interaction with HIV-1 Rev, hence impacting the replication, transcription and translation of the HIV-1 genome in mammalian cells (19–21). eIF5A also mediates the export of *NOS2* mRNA and *TSC2* mRNA in mammalian models (22, 23). Furthermore, eIF5A has been found to be associated with RNA polymerase II (RNA Pol II) in the nuclei of precursor neurons (24). Under hypoxic conditions, the nuclear eIF5A2 isoform binds to the promoter region of the hypoxia-inducible factor 1α (HIF-1α), although its role in HIF-1α activation is not fully clear (25, 26). The eIF5A2 isoform also indirectly regulates the transcription of ageing genes in human neuroblastoma cells by modulating transcription factors associated with the unfolded protein response (27).

Although transcription and translation are considered independent processes due to their distinct molecular mechanisms, timing and sites of action, several factors have been proposed to coordinate them, however, the crosstalk between the two processes is poorly understood (28, 29). Here, we describe a novel function of nuclear yeast eIF5A in transcriptional regulation. By binding to specific coding regions, eIF5A attenuates the recruitment of RNA Pol II to tightly control the expression of the eIF5A translation-dependent mRNAs. Furthermore, we demonstrate that this transcriptional regulation is driven by the eIF5A-dependent motifs, which allow for positive translation regulation in the cytoplasm while driving negative transcriptional regulation in the nucleus, resulting in the fine-tuning of translation efficiencies. These findings expand our understanding of the role of eIF5A in coordinating the various stages of gene expression.

## Results

### Nuclear eIF5A binds to the chromatin of genes that depend on eIF5A for translation

Although eIF5A shuttles between the nucleus and cytoplasm in mammalian and yeast cells, its nuclear functions and putative DNA binding properties still remain unclear. Therefore, we aimed to address whether eIF5A binds to DNA chromatin of specific genes. To answer this question, we performed ChIP-seq experiments in wild-type *Saccharomyces cerevisiae* cells exponentially grown in glucose-based media using a specific eIF5A antibody. Our results revealed that the cross-linking procedure trapped eIF5A on the chromatin and metagene analysis of two biological replicates demonstrated that this binding occurs in transcribed regions (Figure 1A). We examined the functional categories (GOs) enriched in genes showing the strongest eIF5A binding. We found enrichment in categories including “cell wall organization”, “bud growth”, “actin filament organization”, “cytoskeleton organization”, and “filamentous growth” among others (Figure 1B). Interestingly, some of these GOs contain genes whose translation relies on eIF5A as they contain a significant number of eIF5A-dependent motifs such as those with three or more consecutive prolines (9, 10, 30).

**Figure 1.**
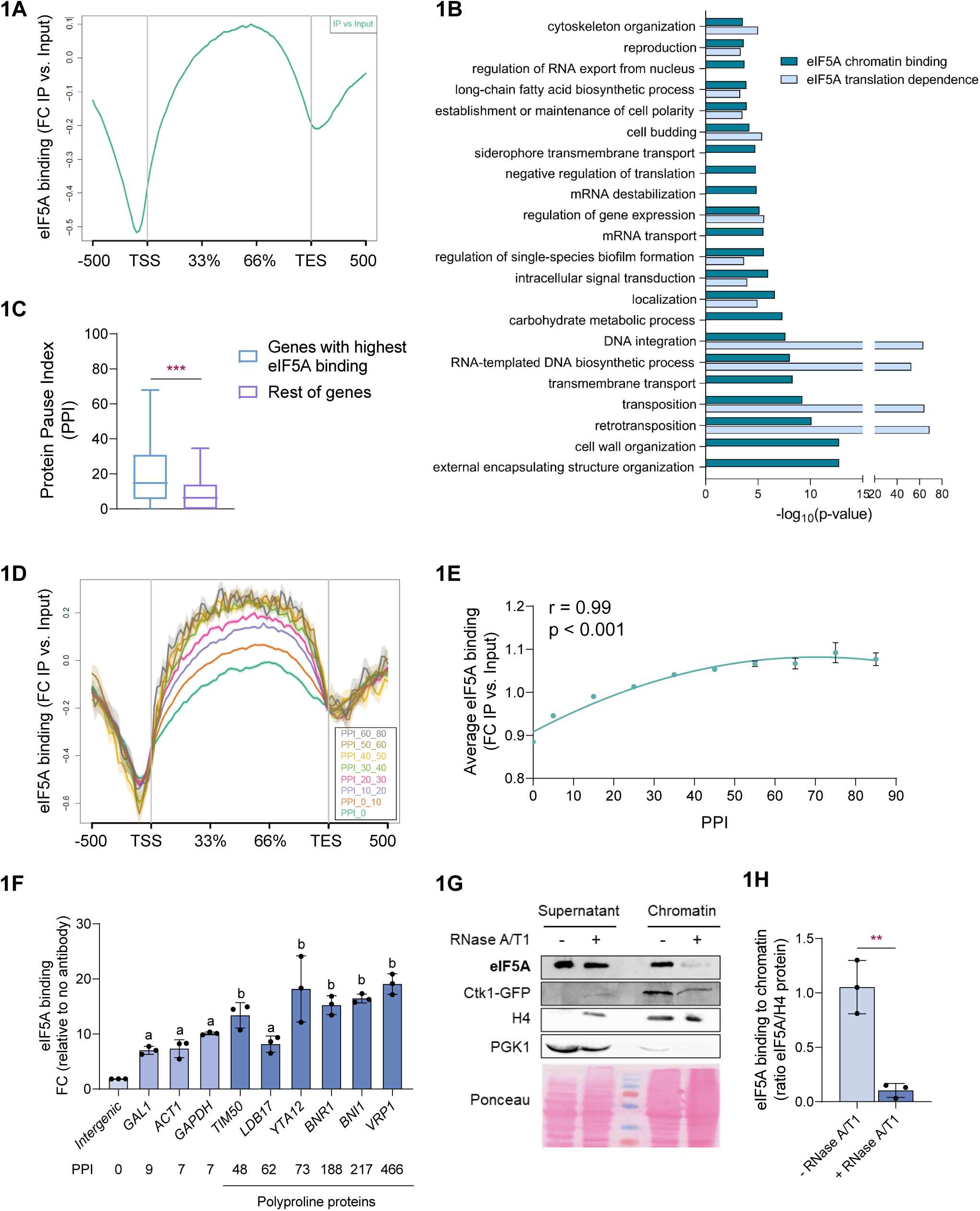
eIF5A binds preferentially to the chromatin of genes whose translation is dependent on eIF5A. (A) Metaplot illustrating the general binding profile of eIF5A to yeast genes obtained by ChIP-seq. Wild-type yeast cells were exponentially-grown in YPD at 25°C and then ChIP-seq of eIF5A was performed using an anti-eIF5A antibody. The graph shows the global eIF5A distribution in the gene body region, from the transcription start site (TTS) to the transcription end site (TES), corrected by the whole cell extract (metaplot of 5143 genes corresponding to two independent ChIP-seq experiments). (B) Biological process Gene Ontology (GO) terms overrepresented in genes with the highest eIF5A binding or the highest eIF5A translation dependence measured with the Protein Pause Index (PPI). Gene set enrichment analysis was done using the ReviGO software. (C) Distribution of the PPI data from the top genes with the highest eIF5A binding or the rest of genes is presented in box-plots, showing the minimum, first quartile, median, third quartile and maximum. Statistical significance was determined using a two-tailed paired Student’s t-test. (D) Metaplot illustrating eIF5A binding distribution in the gene body region corrected by the whole cell extract for genes included in each group with different PPI intervals (see Figure S1A). (E) The average eIF5A binding value relative to input from ChIP-seq analyses is shown for the genes included in each PPI interval group. Experimental data were adjusted to potential trend. The Standard error (SE) and Pearson’s correlation coefficient and the associated significance for the plot is shown. (F) ChIP analysis of eIF5A recruitment in wild-type cells exponentially-grown in YPD at 25°C. ChIP of eIF5A was performed using an anti-eIF5A antibody. The immunoprecipitated DNA was used to quantify the binding to the different genes by qPCR using primers designed for amplification in the specific ORF regions. The percentage of the signal obtained in each ChIP sample with respect to the signal obtained with the DNA from the corresponding whole cell extract was calculated and represented relative to signal of non-antibody sample. Statistical significance was determined using a two-tailed paired Student’s t-test relative to intergenic region-binding signal. When bars do not share any common letter, values are statistically different. (G) Chromatin association of eIF5A depends on RNA. Wild-type yeast cells containing a *CTK1*-GFP genomic fusion and exponentially grown in YPD at 25°C were lysed and then treated or not with an RNase A/T1 mix. After this, chromatin was purified as described in M&M and proteins in the cytoplasmic and chromatin fractions were analyzed by Western blotting using an antibodies against eIF5A, GFP, nuclear histone H4 and cytoplasmic Pgk1 proteins. Ponceau staining is shown as protein loading control. (H) Quantification of eIF5A protein levels, with respect histone H4 protein levels for experiments as in (G). It is shown the mean of eIF5A/H4 ratio of chromatin-associated proteins ± SD from three independent experiments. Statistical significance was determined using a two-tailed paired Student’s t-test of non-treated relative to RNase treatment samples, *p<0.05, **p<0.01, ***p<0.001. n.s means no significant differences.

To shed light into this, and given that the number of experimentally proven eIF5A direct translational targets is relatively low, we classified the yeast genes according to their putative eIF5A-dependence for translation. First, we defined a Protein Pause Index (PPI) for each gene by using the strength pause values obtained for the top 43 eIF5A-dependent tripeptides revealed by 5PSeq analysis (9). The three top tripeptides among these were KPP, PPP and PGW, with pause scores of 8.982, 7.032 and 6.410 respectively (signal of stalled ribosomes under eIF5A depleted conditions vs. non-depleted); while the rest were mostly combinations of proline, glycine and/or acid or basic amino acids (9). The PPI was calculated for each gene as the sum of each of these top 43 eIF5A-dependent tripeptides contained in its corresponding amino acid sequence, multiplied by its specific strength pause value. The higher the PPI for the gene, the more likely that translation of the corresponding mRNA will stall upon eIF5A deficiency. The yeast proteome (data from 6714 proteins) showed an average of 2.9 motifs/protein and an average PPI value of 10.84. A GO search according to the PPI showed that several of the categories with the strongest DNA binding of eIF5A are also functional categories over-represented in those genes with the highest eIF5A requirement for translation (Figure 1B). Moreover, we found a positive and statistically significant difference in PPI values between genes with highest eIF5A binding and the rest of the genome (Figure 1C).

We further analyzed the relationship between eIF5A chromatin binding and their eIF5A-dependence for translation by classifying the entire yeast genome into 10 groups showing different PPI intervals (Figure S1A). Notably, genes with the highest PPI values exhibited higher levels of eIF5A association to their gene body compared to those with the lowest PPI values (Figure 1D). In this line, we observed a progressive increase in eIF5A binding as the PPI value increased until reaching a saturation at PPI values above 60, proportional to the presumed eIF5A-dependence for translation (Figure 1E and Figure S1B).

Consistently with the above ChIP-seq results, we further confirmed the presence of eIF5A at the locus of genes with high PPI by ChIP-qPCR experiments (Figure 1F, Figure S1C). We found a significant binding of eIF5A to all the tested gene ORFs, compared to an intergenic region, where the signal was almost absent as expected from the general profile (Figure 1D). However, the genes with high PPI values, including several confirmed translational targets of eIF5A such as *TIM50* (31), *VRP1* (8), *BNR1* or *BNI1* (30), showed significantly higher levels of eIF5A association compared to those with low PPI values (approximately 2-fold signal).

The association of eIF5A with specific gene bodies could be mediated by DNA or RNA binding. The fact that eIF5A ChIP-seq signals are higher at the 3’ ends of transcribed regions (see Figures 1A and 1D) suggests that eIF5A binding could occur through interaction with the emerging mRNA. To further investigate this, we performed chromatin purification experiments on yeast cells that had been previously lysed and then incubated with a buffer with or without RNases. Proteins associated with chromatin were then detected by Western blotting. Treatment with RNase led to a significant decrease in the levels of chromatin-associated eIF5A (Figure 1G), as well as those of the RNA Pol II serine 2 kinase Ctk1, whose chromatin association is known to be strongly influenced by nascent RNA (32). Conversely, RNase treatment did not interfere with histone H4 binding to DNA, as expected (Figure 1G). The eIF5A/H4 ratio of chromatin-associated proteins was more than 4-fold higher (p-value <0.05) in samples of chromatin purified without RNase pretreatment (Figure 1H).

Altogether, these results confirm that nuclear eIF5A is bound to the ORF of many genes and that eIF5A binds preferentially to the coding regions of genes encoding target mRNAs requiring this factor for their translation. Moreover, eIF5A association to chromatin is highly dependent on RNA.

### Deficiency of eIF5A increases the transcription and mRNA levels of the genes it binds to, and which require it for their translation

The interaction of eIF5A with chromatin suggests that this protein plays a role in transcriptional regulation and mRNA synthesis. To elucidate the potential effects of eIF5A depletion on mRNA synthesis and metabolism, we determined the synthesis rate (SR) and mRNA amount (RA) of the entire yeast genome using the Genomic Run-On (GRO) method (33). We used a wild-type strain and the temperature-sensitive strain *tif51A-1* (carrying a single Pro83 to Ser mutation), since eIF5A is an essential protein (34, 35). The two isogenic yeast strains were grown in glucose-based medium to early exponential phase at permissive temperature (25°C) and then transferred to restrictive temperature (37°C) for 4 hours. At this point, eIF5A protein levels were almost undetectable in *tif51A-1* cells at 37°C (Figure S2A), and the growth rate was decreased (Figure S2B,C), without affecting mutant cells viability (30).

We first analyzed all the yeast protein-coding genes (4,505 genes) and found that eIF5A depletion reduced global transcription (Figure 2A left), as expected for a reduction in growth rate (36), but increased total RNA (Figure 2D left). mRNA half-life estimations revealed a significant stabilization of most transcripts in *tif51A-1* cells, indicating that global mRNA stabilization is the main contribution to the increase in RA under eIF5A deficiency.

**Figure 2.**
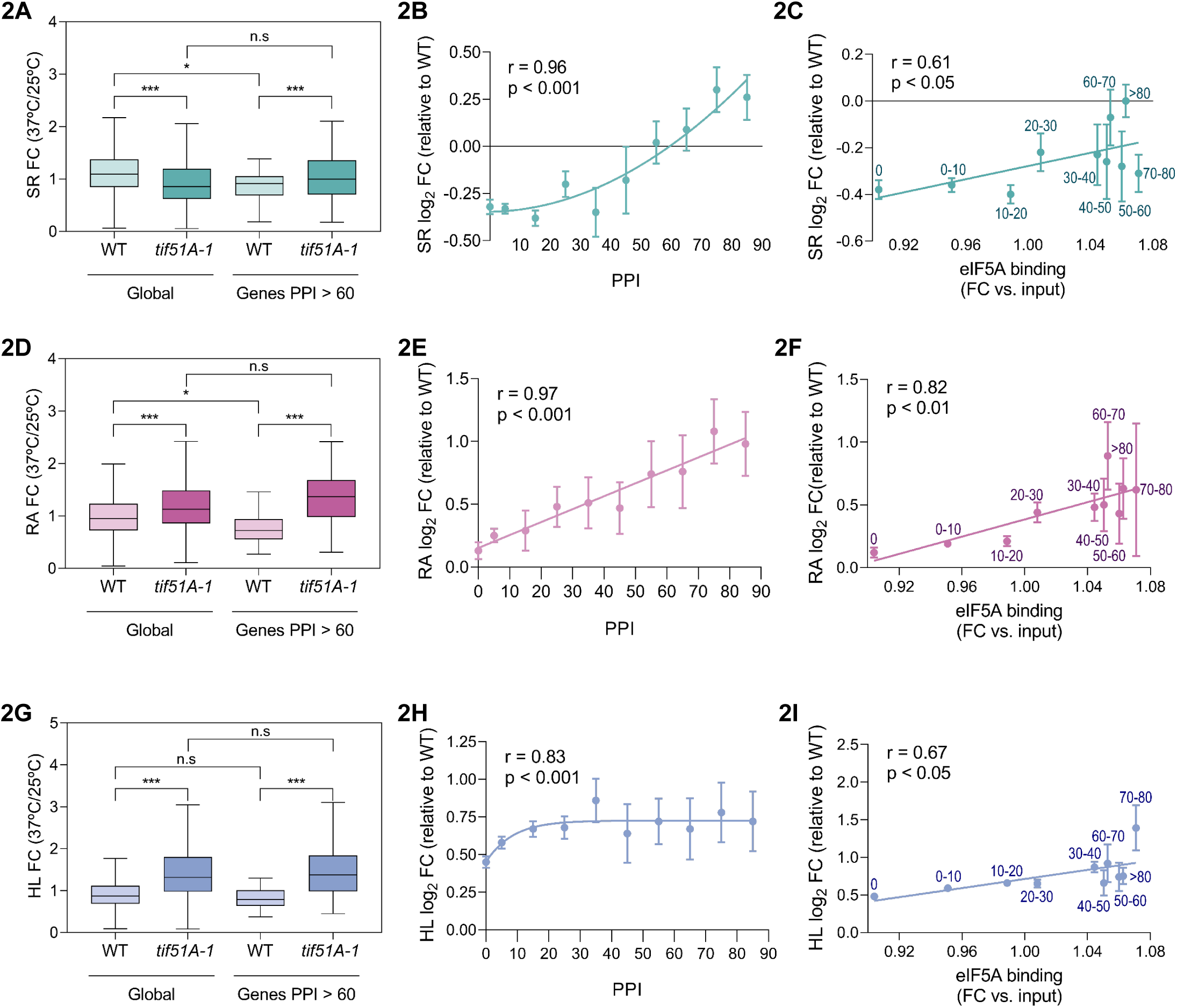
Changes in synthesis rate (SR) and RNA amount (RA) upon eIF5A depletion correlate with the degree of eIF5A-dependency for translation and with the binding of eIF5A to chromatin. Wild-type and *tif51A-1* mutant yeast strains were grown in YPD medium at 25°C until early exponential phase and then transferred to 25°C and 37°C for 4 hours. Then cells were collected and processed as described in M&M for GRO assay. (A,D,G) Genomic data of SR (A), RA (D) and mRNA half-lives (HL) (G) are presented in box-plots, showing the minimum, first quartile, median, third quartile and maximum from the global data (4505 genes) and from genes with PPI > 60 (48 genes). Data are presented as the fold change (FC) 37°C *vs.* 25°C from three independent experiments. Statistical significance was determined using a two-tailed paired Student’s t-test relative to corresponding wild-type cells. ***p<0.001. (B,E,H) Graphs represent the average SR (B), RA (E), and HL (H) for all genes included in each Protein Pause Index (PPI) interval group. Values are given as the log_2_ fold change (FC) at 37°C vs. 25°C ± S.E of *tif51A-1* strain, relative to the corresponding value in the wild-type strain. Experimental data were adjusted to logarithmic (B), linear (E) or potential (H) trends. (C,F,I) Graphs represent the average SR (C), RA (F), and HL (I) for all genes included in different PPI group (the labels indicate the PPI interval) versus the average eIF5A binding from ChIP-seq analyses for each group of genes. Values represent the log_2_ fold change (FC) at 37°C vs. 25°C ± S.E of *tif51A-1* strain, relative to the corresponding value in the wild-type strain. Experimental data were adjusted to linear trends. (B,C,E,F,H,I) Pearson’s correlation coefficient and the associated significance for the plots are shown.

To study the changes occurring in the set of eIF5A translation-dependent genes, we established a target group of genes in which their PPI was above 60 (48 genes). These genes were strong candidates to be dependent on eIF5A for their translation and included most of the yeast eIF5A-identified targets (Figure S1C). We observed that the synthesis rate (SR) of these genes with high eIF5A-dependence had a significant increase upon eIF5A depletion, opposite to the decreased SR observed globally (Figure 2A). The analysis of the RA among this group of genes showed a more marked and significant RA increase than the global data (Figure 2D), whereas the analysis of the HL showed minor differences with regard to the global data (Figure 2G). Therefore, we identified specific trends for genes predicted to behave as eIF5A translational-dependent, which diverged substantially from the general trend in the *tif51A-1* mutant.

To further investigate this, we used the previous classification of genes according to their PPI values (Figure S1A). We calculated the relative fold change 37°C vs. 25°C for each gene in the *tif51A-1* mutant respect to this fold change in the wild-type, and then the mean values of SR, RA and HL and plotted them against the PPI values and the corresponding eIF5A binding values from ChIP-seq for each PPI gene group. These analyses revealed that the relative changes of SR in the eIF5A mutant correlated significantly and positively with the PPI and with the eIF5A binding to chromatin (Figures 2B,C and S2D-G). Consistent with this increase, we observed a proportional increase in RA that was strongly and positively correlated with both the PPI and the eIF5A binding (Figures 2E,F and S2H-K). In contrast, the HL data showed a lower correlation with the PPI and with the eIF5A binding to chromatin (Figures 2H,I and S2L-O).

As eIF5A association with chromatin depends largely on RNA (Figure 1G), we analyzed whether eIF5A binding correlates with gene transcription levels in wild-type or with changes in gene transcription that may occur when cells are shifted from 25°C to 37°C to deplete eIF5A protein. We found a modest positive correlation between eIF5A binding to gene bodies and transcription levels (Figure S2P) and a very slight correlation between eIF5A binding and SR change during the temperature shift (Figure S2Q). Together with our previous results (Figure S1C), these observations suggest that the recruitment of eIF5A to chromatin is largely influenced by PPI, and slightly dependent on the synthesis rate.

In sum, our genomic studies concluded that eIF5A depletion yields overall mRNA increase in yeast cells because of global mRNA stabilization, while the levels of mRNAs with both stronger eIF5A-dependence for their translation and eIF5A binding to their chromatin increase due to a higher synthesis rate rather than stability. Therefore, these findings suggest a specific role of nuclear eIF5A in the transcriptional repression of genes encoding its target proteins.

### Nuclear localization via an N-terminal NLS and cytoplasmic hypusination of eIF5A are critical for transcriptional repression of eIF5A translation-dependent genes

To validate our genome-wide data, we measured the changes in mRNA levels (RA) of several specific genes showing differences in their eIF5A translation-dependencies, according to their PPI values. Results showed a statistically significant increase in mRNA levels in eIF5A-depleted *tif51A-1* cells with respect to the wild-type for most of the tested genes (Figure 3A). Consistent with our GRO results, this increase correlated well with the PPI values, indicating that the stronger the eIF5A translation dependency, the higher the mRNA abundance.

**Figure 3.**
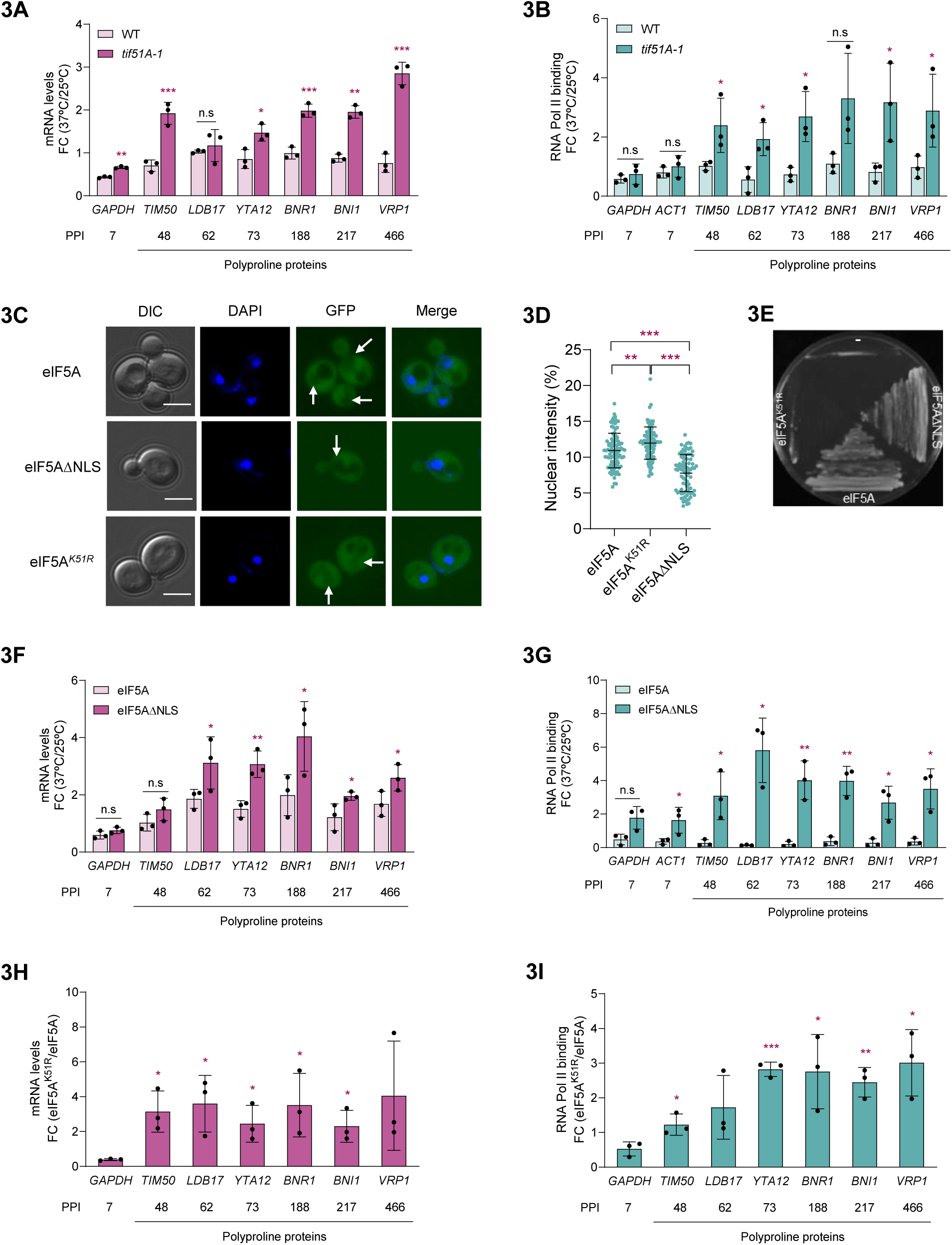
eIF5A depletion, nuclear exclusion, or deficient hypusination increases the transcription of eIF5A translation-dependent genes. (A,B) Wild-type and *tif51A-1* mutant yeast strains were grown in YPD medium at 25°C until exponential phase and then transferred to 25°C and 37°C for four hours. (A) mRNA levels from each gene were determined by RT-qPCR using primers designed for amplification in the ORF regions. (B) ChIP of RNA Pol II was performed using the antibody anti-Rpb1 (8WG16). The immunoprecipitated DNA was used to quantify the binding to the different genes by qPCR using primers designed for amplification in the ORF regions. The percentage of the signal obtained in each ChIP sample with respect to the signal obtained with the DNA from the corresponding whole cell extract was calculated. (C) Wild-type strain expressing a second copy of GFP-eIF5A, GFP-eIF5AΔNLS or GFP-eIF5A*^K51R^* were cultured in YPD medium until reaching exponential phase and subjected to fluorescence microscopy. Cells were incubated for 5 min with DAPI prior microscopy to stain the nuclei. White arrows indicate the nuclei. A representative image is shown from three independent experiments. Scale bar, 4 μm. (D) Quantification of percentage of nuclear signal is shown from a minimum of 100 cells. Results are presented as individual values together with the mean ± SD from three independent experiments. The statistical significance was measured by using a two-tailed paired Student t-test. (E) *tif51A-1* strain with or without (-) the expression of a second copy GFP-eIF5A, GFP-eIF5AΔNLS or GFP-eIF5A*^K51R^* was cultured in YPD at 37°C to deplete eIF5A from the first copy (tif51A-1 protein). (F,G) Wild-type expressing GFP-eIF5A or GFP-eIF5AΔNLS in the *TIF51A* locus were grown as in (A) and mRNA levels (F) and RNA Pol II ChIP binding (G) were measured as in (A,B). (A,B,F,G) Data are presented as the mean fold change 37°C vs. 25°C ± SD from three independent experiments. Statistical significance was determined using a two-tailed paired Student’s t-test relative to wild-type cells. (H,I) Yeast strain with tetO_7_-*TIF51A* at the endogenous locus and a second copy of GFP-eIF5A or GFP-eIF5A*^K51R^* were cultured overnight in YPD at 25°C in the presence of doxycycline (15 µM) to deplete eIF5A from the first copy, and then mRNA levels (H) and RNA Pol II ChIP binding (I) were measured as in (A,B). Data are presented as the mean fold change eIF5A*^K51R^* vs. eIF5A wild-type ± SD from three independent experiments. Statistical significance was determined using a two-tailed paired Student’s t-test relative to wild-type cells. *p<0.05, **p<0.01, ***p<0.001. n.s means no significant differences.

To independently corroborate that the increase in mRNA levels of eIF5A-translation target genes upon eIF5A depletion is due to an increase in SR (Figures 2D-F), we investigated the occupancy of total RNA Pol II, as a proxy of transcriptional activity. We performed chromatin immunoprecipitation (ChIP) using an antibody against the catalytical Rpb1 subunit of RNA Pol II and observed that total RNA Pol II was recruited to similar levels in wild-type and *tif51A-1* cells to eIF5A translation-independent genes, such as *ACT1* or *GAPDH*. However, and consistent to our GRO results, we found a higher and significant association (2- to 4-fold increase) of the total RNA Pol II to the ORFs of genes showing a strong eIF5A-dependence for their translation in the *tif51A-1* cells (Figure 3B). The higher binding of the RNA Pol II elongation complex was in accordance to the higher mRNA levels of the affected genes (Figure 3A).

We next investigated whether the transcriptional derepression observed upon eIF5A depletion was linked to a nuclear function of eIF5A. To address this, we first identified a nuclear localization signal (NLS) in the eIF5A yeast protein in order to split nuclear and cytoplasmic eIF5A functions. In mammalian cells it has been proposed that the eIF5A N-terminal protein region acts as NLS, although bearing no structural similarity with classical nuclear localization signals (37). In *S. cerevisiae*, the N-terminal (1-MSDEEHTFETADAGSSATY-19) extension of the eIF5A protein is enriched in acidic residues and conserved across eukaryotes but absent in bacteria and archaea orthologues (Figure S3A) and, therefore, candidate to act as NLS. We constructed yeast strains expressing an additional copy of full eIF5A sequence or a ΔNLS version of eIF5A (Δ1-19) fused to GFP. Importantly, both GFP-tagged eIF5A versions demonstrated to be functional (Figure 3E). Direct *in vivo* visualization of GFP-eIF5A revealed a whole cell distribution pattern with a primary cytoplasmic signal, consistent with its cytoplasmic function, and a faint nuclear signal corresponding to approximately 12% of the total signal (Figures 3C,D). This partial nuclear localization of eIF5A under standard conditions aligns well with previous observations in mammalian cells (13, 17, 18, 37–39) and confirms the shuttling of eIF5A between nucleus and cytoplasm. However, deletion of the N-terminal region in yeast eIF5A (GFP-eIF5AΔNLS) hindered its nuclear import (Figures 3C,D). This demonstrated that the N-terminal NLS is necessary for signaling eIF5A nuclear accumulation in *S. cerevisiae* cells. Conversely, deletion of Pdr6, the described nuclear exportin of eIF5A (14, 15), resulted in increased nuclear accumulation of GFP-eIF5A (Figures S3B,C). Interestingly, the nuclear localization of eIF5A was found to be specific and clearly different from that of other translation factors such as eIF4e, which is clearly excluded from the nucleus under standard conditions (Figures S3B,C).

Because both GFP-eIF5A and GFP-eIF5AΔNLS as second copy complemented growth of *tif51A-1* mutant at 37°C (Figure 3E), we generated new strains with either of the two eIF5A versions at its natural chromosomal locus and confirmed their viability (Figure S3D). As expected, blocking the nuclear import of eIF5A compromised its association to the chromatin (Figure S3E). Then, we examined the changes in mRNA levels of genes with different eIF5A translation-dependencies (Figure 3F) and found that cells with the eIF5AΔNLS version exhibited a statistically significant increase in mRNA levels compared to wild-type cells, and that this increase correlated with the PPI values (Figure 3F). In parallel, we tested the association of RNA Pol II by ChIP and observed a significantly higher recruitment (3- to 5-fold increase) to the ORFs of genes with the highest PPI values (Figure 3G). Importantly, changes in both mRNA levels and RNA Pol II binding in eIF5AΔNLS cells were similar to those in *tif51A-1* cells.

We then explored whether inhibiting the translational activity of eIF5A might affect its nuclear function in gene repression. To achieve this, we used an eIF5A version with a hypusine site mutation (K51R), which cannot be modified into the hypusine form (40). First, we confirmed in yeast cells carrying GFP-eIF5A^K51R^ as second copy that the mutation induced eIF5A nuclear accumulation and loss of protein function (Figures 3C,D,E), consistent with previous reports (17, 39). Next, we constructed a yeast strain in which the *TIF51A* gene (expressing eIF5A), was under the control of a tetO_7_ promoter and confirmed that switching off the native eIF5A gene resulted in lack of growth and increased mRNA levels of the eIF5A translation-target genes, as it happens with the *tif51A-1* temperature-sensitive mutant at restrictive temperature (Figures S3F,G). Then, we engineered this conditional yeast strain to add either GFP-eIF5A^K51R^ or a control GFP-eIF5A version as a second copy. As expected, both the GFP-eIF5A^K51R^ and the GFP-eIF5A proteins were expressed, but only the second version was hypusinated and rescued growth after the addition of doxycycline (Figures S3G,H,I). After doxycycline addition, we observed that both the mRNA levels and RNA Pol II recruitment increased for the genes encoding translational targets of eIF5A in cells expressing the eIF5A^K51R^ mutant protein respect to the ones expressing GFP-eIF5A (Figures 3H,I). The changes in both mRNA levels and RNA Pol II binding in eIF5A^K51R^ cells were similar to those in *tif51A-1* cells.

Taken together, these results suggest that the repression of transcription by eIF5A requires its nuclear localization, as this is abolished when its NLS is removed. Moreover, it also requires a functional eIF5A for cytoplasmic translation, since the non-hypusinated eIF5A protein, which cannot bind ribosomes, is unable to repress transcription despite presenting a higher nuclear accumulation. Therefore, these results suggest that nuclear eIF5A plays a direct role in transcriptional repression and that this is functionally linked to its activity in translation.

### The presence of polyproline motifs in eIF5A translation-dependent genes is sufficient to promote their repression by eIF5A

Our discovery that eIF5A preferentially binds to the chromatin of genes that encode target proteins for translation –a process that is predominantly RNA-mediated- and its correlation to the transcriptional regulation by eIF5A, prompted us to investigate the chromatin/RNA feature that mediates eIF5A specificity. Several of the strongest identified eIF5A translation-targets contain polyproline motifs, resulting in high PPI values (9, 10). We explored the effect in transcription of removing these motifs from eIF5A translation targets and of adding these motifs to translation-independent genes.

Following this idea, we first obtained yeast strains targeting the genes *YTA12*, *TIM50* and *BNR1*, which have high PPI values and have been previously shown to be bona-fide translation targets of eIF5A (30, 31, 41). Here, we have shown that these genes present high levels of eIF5A binding (Figure 1F), and that respond to nuclear eIF5A depletion by increasing SR and mRNA levels (Figures 3A,B,F,G). For each gene, we obtained two genomic GFP- or HA-tagged versions in wild-type and *tif51A-1* cells: one with GFP or HA after the native coding region (*YTA12*-GFP, *BNR1*-HA and *TIM50*-GFP) and a mutated version in which the region containing the polyPro stretches was deleted (*YTA12ΔPro*-GFP, *BNR1ΔPro*-HA and *TIM50ΔPro*-GFP). These new versions, therefore, showed lower PPI values and were predicted to behave as eIF5A translation-independent genes (Figures 4A and S4A,B; (30).

**Figure 4.**
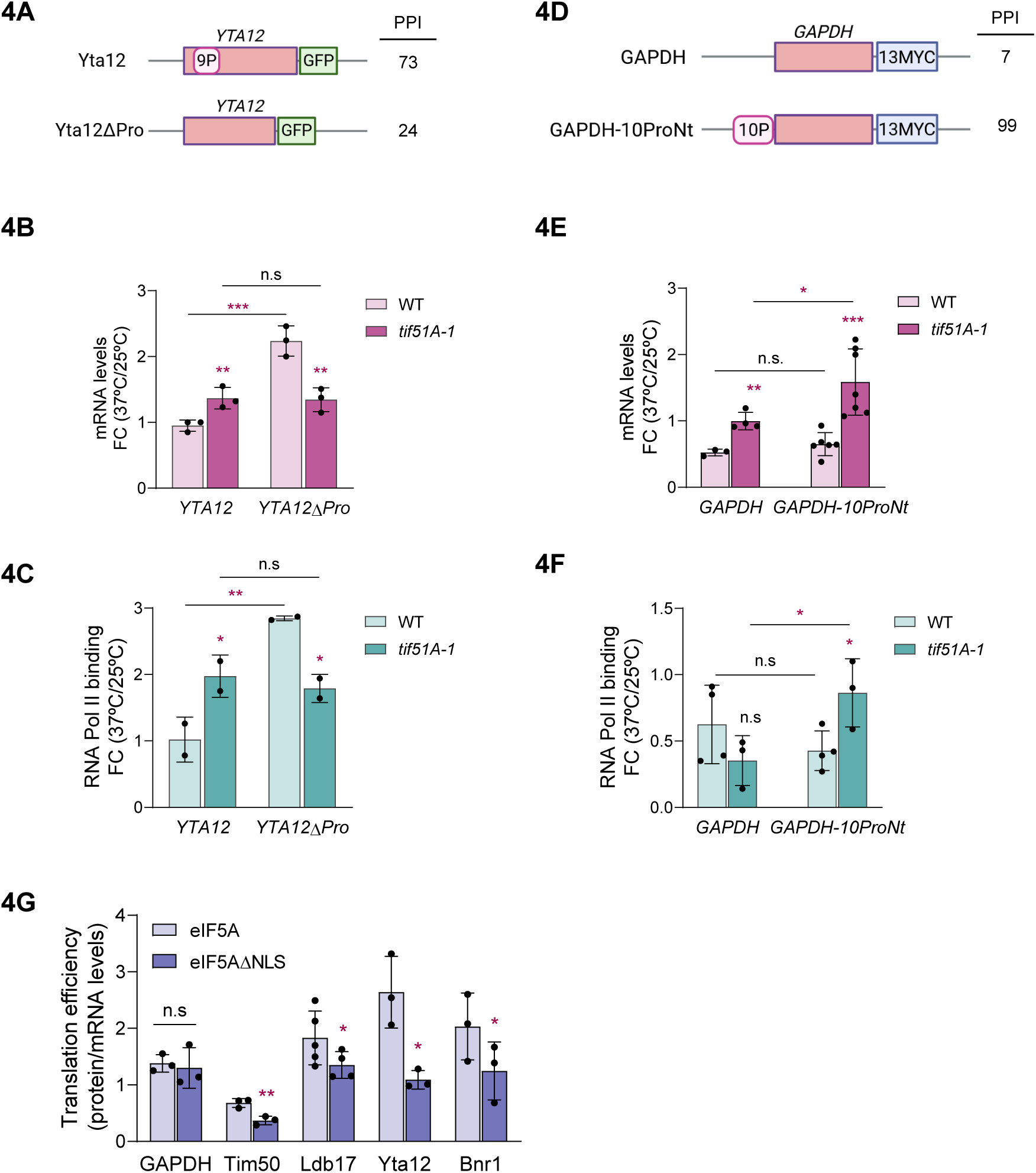
Transcriptional repression driven by polyPro encoding sequences promotes the translation efficiency of eIF5A target proteins. (A,D) Schematic diagram showing the C-terminal genomic tagging of native *YTA12* (A) and *GAPDH* (D) ORFs as well as their polyPro-deleted (A, *YTA12ΔPro*) or -inserted (D, *GAPDH*-10ProNt) versions. PPI of all gene versions are shown. (B,E) Wild-type and *tif51A-1* mutant yeast strains carrying the native and the mutated versions of *YTA12* and *GAPDH* genes were grown in YPD medium at 25°C until exponential phase and then transferred to 25°C and 37°C for four hours. mRNA levels from *YTA12* (B) and *GAPDH* (E) versions were determined by RT-qPCR. (C,F) Wild-type and *tif51A-1* mutant yeast strains carrying the native and the mutated versions were grown as in (B,E). ChIP of RNA Pol II was performed using the antibody anti-Rpb1 (8WG16). The immunoprecipitated DNA was used to quantify the binding to *YTA12* (C) and *GAPDH* (F) genes by qPCR using primers designed for amplification in the specific ORF regions. The percentage of the signal obtained in each ChIP sample with respect to the signal obtained with the DNA from the corresponding whole cell extract was calculated. (G) Wild-type cells expressing the native eIF5A or the eIF5AΔNLS version were grown in YPD medium at 30°C until exponential phase. mRNA and protein levels from *GAPDH*, *TIM50*, *LDB17*, *YTA12* and *BNR1* genes were determined and quantified (see supplemental Figure S5), and then translation efficiency was determined as the ratio between protein and mRNA levels. (B,C,E,F,G) Data are presented as the mean ± SD from at least three independent experiments. Statistical significance was determined using a two-tailed paired Student’s t-test relative to corresponding wild-type cells. *p<0.05, **p<0.01, ***p<0.001. n.s means no significant differences.

We analyzed the mRNA levels in the new strains and found similar results for the three genes. We found a significant increase in the mRNA levels of the three native genes (expressing the polyPro motifs) in the *tif51A-1* cells following the temperature shift. However, we found that this mRNA increase was no longer present under eIF5A depletion for the mutated versions of the three genes without the polyPro motifs (Figures 4B and S4C,D). Moreover, we observed that the polyPro sequences had a repressive effect in the presence of eIF5A (wild-type strain), as the mRNA levels were lower for the *YTA12*, *BNR1* and *TIM50* genes carrying the polyPro motifs than for the ΔPro protein versions (Figure 4B and Figure S4C,D). Next, we evaluated the recruitment of the RNA Pol II to the *YTA12* gene by performing ChIP analysis of RNA Pol II. We found that polymerase binding to the *YTA12* gene body was higher upon eIF5A depletion in the *tif51A-1* cells, but only for the polyPro-containing *YTA12*-GFP gene and not for the YTA12ΔPro-GFP gene without the proline stretches (Figure 4C).

Next, we aimed to test the impact on mRNA levels and transcription following the insertion of a polyPro region into an eIF5A-translation independent gene. For this, we generated wild-type and *tif51A-1* strains targeting the *GAPDH* gene (isoform *TDH1*), that encodes the glyceraldehyde-3-phosphate dehydrogenase enzyme involved in glycolysis and gluconeogenesis. We generated genomic C-terminal Myc-tagged strains: one with the native coding region (*GAPDH*-myc) and low PPI value, and a mutated version in which 10 consecutive prolines inserted at the N-terminal region after the first codon (GAPDH-10ProNt-myc). As expected, only the version with polyPro motifs (high PPI) behaved as an eIF5A-dependent gene for translation (Figures 4D and S4E,F).

Using these new strains, we observed a slight but significant increase in the mRNA levels of the native gene (*GAPDH*-myc) in the *tif51A-1* cells after the temperature shift. However, we found a significantly greater increase in the mRNA levels of the *GAPDH*-10ProNt-myc version in the *tif51A-1* cells (Figure 4E). We also observed that the association of the RNA Pol II to the *GAPDH* gene body was similar across strains carrying the *GAPDH*-myc whereas its recruitment was significantly higher in *tif51A-1* cells carrying the GAPDH-10ProNt-myc version compared to the wild-type (Figure 4F).

In summary, these results suggest that the presence of polyPro motifs, which stall translation under eIF5A deficiency, are both necessary and sufficient to drive the eIF5A-dependent transcriptional regulation.

### The coordinated nuclear and cytoplasmic functions of eIF5A facilitate the efficient synthesis of its target proteins

We were prompted to investigate the biological advantages of this double but opposite eIF5A-dependent gene regulation, as the results suggested that nuclear eIF5A drives transcriptional repression while promoting the cytoplasmic translation of the same targets. To do this, we used the eIF5AΔNLS version, which is mostly excluded from the nucleus (Figures 3C,D), does not repress transcription (Figures 3F,G) but retains its translation function, thereby maintaining cellular viability (Figure 3E). We then compared this ΔNLS version with the native eIF5A version, which is active in both the nucleus and the cytoplasm. As previously demonstrated, both eIF5AΔNLS and eIF5A strains grew well in glucose media at 30°C and had similar doubling times (Figure S5C); therefore, this result did not indicate superior performance of native eIF5A under standard conditions. We then investigated whether translation efficiency differed in yeast cells carrying either of the eIF5A versions. To do this, we measured the mRNA and protein levels of several eIF5A translation-dependent genes (*TIM50*, *LDB17*, *YTA12* and *BNR1*), as well as a control gene (*GAPDH*), and then calculated the protein/mRNA ratio as an indicator of translation efficiency. The results again showed that inhibiting the nuclear-cytoplasmic shuttling of eIF5A increased the mRNA levels of target genes (Figure S5D); however, we did not observe an increase in the corresponding protein levels (Figures S5E,F). Consequently, estimation of the protein-to-mRNA ratios revealed lower translation efficiencies for eIF5A-dependent genes when nuclear localization of eIF5A was inhibited (Figure 4G), which was not due to lower eIF5AΔNLS expression (Figures S5A,B). Thus, these results suggest that reducing transcription by nuclear eIF5A positively affects the subsequent eIF5A-dependent cytoplasmic translation of the corresponding target mRNA. However, this slight difference in the translation efficiency of target genes does not result in improved growth under normal conditions.

## Discussion

In recent years, the crosstalk between mRNA synthesis, maturation, nuclear export and decay has been established (28). However, there is less information on the interaction between two of the most physically and temporally separated processes: transcription and translation.

Here, we found that the highly abundant translation elongation factor eIF5A associates with the chromatin of many genes in yeast cells. However, we identified a group of genes with higher binding of eIF5A where eIF5A appears to act as a repressor, indicated by an increase in transcription, RNA Pol II binding and the corresponding mRNA levels upon eIF5A depletion, and showing a different pattern to global regulation. This group of genes mainly encodes proteins with high levels of tripeptide motifs (e.g. polyPro), which have been identified as conferring dependency on eIF5A for translation of the corresponding mRNA (9, 10). While there is currently no experimental evidence for the dependency on eIF5A for the translation of all genes in this group with high PPI, we have shown that several of them are bona fide eIF5A translation targets (e.g. *TIM50*, *YTA12*, *BNI1*, *BNR1*, *VRP1*), that are bound by eIF5A, and that their RNA Pol II association and mRNA levels depend on the presence of nuclear eIF5A. These results strongly suggest a direct role for eIF5A in negatively regulating their transcription.

Our results point that transcription is regulated by the presence of eIF5A in the nucleus, since the regulation is lost when using a version of eIF5A without the identified nuclear localization signal. However, the precise mechanism involved still remains unknown. Previous data have shown that eIF5A interacts with RNA Pol II in precursor neurons (24), and that under hypoxia the mammalian eIF5A2 isoform binds to the HIF1-1a gene where it may positively regulate its transcription (25, 26). The presence of a conserved NLS in the N-terminal extension of the eukaryotic eIF5A proteins suggests that nuclear localization of the protein has a relevant function that has been conserved through evolution. Beyond its nuclear localization, this transcriptional regulation also requires the active engagement of cytosolic eIF5A in translation, as it is lost in cells expressing the eIF5A^K51R^ mutant, which cannot undergo hypusination and therefore fails to bind ribosomes and regulate translation. This result suggests a link between the two activities of the protein.

Furthermore, our results point to eIF5A’s association with chromatin relying on the presence of RNA. This suggests that, like other RBP proteins, eIF5A may bind to nascent RNA co-transcriptionally to modulate transcription. eIF5A structural features suggest a potential to interact with nucleic acids. The C-terminal domain resembles the cold-shock domain, common in DNA and RNA-binding proteins, while the N-terminal carries the hypusine residue, which contains two positive charges and resembles spermidine, a molecule known to interact with DNA or RNA (42, 43). In this line, eIF5A has been previously described to interact directly or indirectly with mRNAs (20, 44). For instance, it functions as a co-factor of the HIV-1 Rev protein, thereby modulating transcription, translation, and replication of the viral genome in mammalian cells (19, 21). Beyond viral infection, eIF5A has been linked to mRNA export, including *NOS2* in a diabetic inflammation model (22) and *TSC2* under anaerobic stress (23). Recently, eIF5A was identified as an RNA-binding protein in a time-resolved profiling study in mammalian cells, where it was unexpectedly captured bound to mRNAs at early stage points of mRNA life cycle and before nuclear export (45).

As the binding of eIF5A to nascent mRNA seems plausible, how it specifically recognizes its target mRNA is a primordial question. We found that nucleotide sequences encoding polyPro motifs confer dependency on eIF5A not only for translation but also for transcriptional regulation. Codons encoding prolines (CCX sequence) create C-rich RNA regions that could form a recognition motif. Similarly, other eIF5A-dependent tripeptide motifs contributing to PPI also contain proline in their sequences (9, 10). In this sense, the presence of specific C-rich motifs in mRNAs has been shown to serve as platform for binding different RBP (46, 47). Further research is needed to clarify this issue.

Our results raise another relevant question: what are the biological benefits of regulating translation and transcription by the same protein, thus yielding coordinated regulation? We demonstrated that the translation efficiency of target genes is reduced when eIF5A is predominantly excluded from the nucleus. Thus, inhibiting the transcriptional repression of eIF5A translation-target genes using eIF5AΔNLS increased mRNA levels, but not the levels of the corresponding proteins. These results suggest that translation efficiency improves when eIF5A associates with chromatin early on. In this context, it has been described that the highly abundant translation elongation factor 1A (eEF1A), essential for delivering aminoacyl-tRNA to the ribosome for translation, also shuttles between the nucleus and the cytoplasm, and binds to the chaperone mRNA encoding Hsp70 to couple its transcription and translation during heat stress (HS) in mammalian cells (48). Altogether, results suggest the possibility of nascent mRNA imprinting by translation elongation factors that shuttle between the nucleus and the cytoplasm, facilitating the subsequent synthesis of encoded proteins. In the case of eIF5A, however, transcription and translation are regulated in opposite directions: transcription is repressed and translation is promoted. We propose that it may be beneficial for the cell to synthesize fewer eIF5A target mRNAs, as the depletion of too much cellular eIF5A protein could interfere with other essential functions. In this respect, recent work has shown that the aggregates of the pathological Huntingtin (mHTT) protein traps eIF5A protein in the brains of mice, and has proposed that this eIF5A depletion disrupts homeostatic control and impairs cellular recovery from stress (49). Additionally, it has recently been proposed that eIF5A could play a role in global translation progression and connect ribosomal progress with quality surveillance (50). Unresolved ribosome stalling in mRNAs due to eIF5A protein scarcity could lead to intense competition for ribosomes, which is the main factor influencing translation costs and, consequently, cellular fitness (51). It is worth noting that the nucleus-cytoplasm shuttling of eIF5A is regulated by post-translational modifications of the protein, such as hypusination, sumoylation, and acetylation (17, 18, 39), which may respond to signaling cues activated by changing conditions. In this line, several studies in mammals have described the translocation of eIF5A to the nucleus in response to hypoxia and apoptosis induction (25, 52). Future systematic studies under different conditions may reveal the beneficial effects of transcription-translation crosstalk by eIF5A.

## Materials and methods

### Yeast strains, plasmids, growth conditions and materials

*S. cerevisiae* strains and plasmids used are listed in Supplementary Tables S1 and S2 respectively. For all the experiments carried out *S. cerevisiae* cells were grown in liquid YPD (2% glucose, 2% peptone, 1% yeast extract).

For detailed procedures of plasmids and strains generation and all materials used see SI Appendix.

### RT-qPCR analysis

For the analysis of the mRNA levels, total RNAs were isolated from yeast cells following the protocol described in (53) The reverse transcription and quantitative PCR reactions were performed as detailed in (54). Endogenous *ACT1* mRNA levels were used for normalization. At least three biological replicates of each sample were analyzed, and the specific primers designed to amplify gene fragments of interest are listed in Table S3. More details can be found in SI Appendix.

### Western blotting

For yeast protein content analysis by western blotting we followed the protocol described in (55). Samples were run on SDS-PAGE gels, transferred, and immunoprobed as described in SI Appendix 1. Antibodies used in this study are listed in the SI Appendix. In order to capture variation across all samples, the signal in each lane was normalized to the mean signal across all lanes in a single blot. Then, the resulting signal of bands was normalized against the corresponding G6PDH resulting signal. At least three biological replicates of each sample were analyzed.

### Fluorescence microscopy and analysis

Yeast cells grown to a logarithmic phase in YPD medium were subjected to standard fluorescence and phase contrast microscopy. Fluorescence images were acquired using an Axio Imager Z1 fluorescence motorized microscope (Carl Zeiss, Germany). Images were recorded with an AxioCam MRm digital camera (Carl Zeiss, Germany). 4′,6-Diamidino-2-phenylindole dihydrochloride (DAPI) was used to visualize nuclei. The same exposure times were used to acquire all images and at least three biological replicates of each sample were analyzed. All the imaging analysis was performed on Image J software. At least 100 single cells were scored from three independent experiments. More detailed procedures can be found in SI Appendix.

### Chromatin immunoprecipitation

The chromatin immunoprecipitation (ChIP) experiments were performed as previously described (56) with the modifications described in (53). qPCR was run as described above using the primers listed in Supplementary Table S3. The qPCR amplification data were normalized with the total input DNA value in the corresponding whole cell extract. Immunoprecipitations of eIF5A were made with anti-eIF5A antibody and Dynabeads anti-rabbit IgG; and of RNA Pol II with anti-Rpb1 antibody and Dynabeads Pan Mouse IgG. At least three biological replicates of each sample were analyzed. More detailed procedures can be found in SI Appendix.

### ChIP-seq and sequencing analysis

Wild-type yeast cells exponentially grown in YPD at 25°C were used for ChIP-seq experiments. Libraries for ChIP-Seq were prepared at IRB Barcelona Functional Genomics Core Facility and submitted for single end 50 nt sequencing on a NextSeq2000 (Illumina). More than 4 Gbp of reads were produced, with a minimum of 17 million of single end reads per sample.

Data analyses was performed at the Statistical and Omics Data Analyses facility of the SCSIE-Universitat de València following the pipeline described in SI Appendix.

Biological process Gene Ontology (GO) terms overrepresented in genes with the highest eIF5A binding or the highest eIF5A translation dependence measured with the Protein Pause Index (PPI) were identified using Gorilla tool and the resulting list of GO identifiers with their adjusted p-values was submitted to ReviGO software. ReviGO was run with the default semantic similarity threshold to cluster redundant GO terms and return a representative subset.

More detailed procedures can be found in SI Appendix.

### Protein Pause Index (PPI) calculation

The translation dependency of each yeast gene on eIF5A was estimated by calculating the PPI. First, the number of the top 43 eIF5A-dependent tripeptide motifs present in the encoded protein amino acid sequence was determined. These motifs cause ribosome pausing when eIF5A is depleted in yeast cells, as described by (9). Secondly, the number of each motif was multiplied by its pause strength value, as revealed by 5PSeq analysis (9). Thirdly, the PPI for each gene was obtained as the sum of motifs x strength.

### Determination of individual transcription rates and mRNA levels

To determine the transcription rate in the corresponding strains, a genomic run-on (GRO) was performed as originally described in (33) and modifications made in (57) Briefly, to perform the run-on, on aliquot of each cell culture under the required conditions was resuspended in the appropriate buffer with [α-^33^P]-UTP and incubated at 30°C for 5 min to allow transcription elongation. Then, radioactive incorporation into nascent mRNA for each yeast gene was measured using home-made macroarray nylon filters (33).

A second aliquot of the same cell culture was used to isolate total RNA and reverse transcribed in the presence of [α-^33^P]-dCTP and, then, labelled cDNA was used to hybridize macroarray filters. Both GRO and cDNA labelled samples belonging to the same sampling were successively hybridized against the same filter. The scanned microarray images from both GRO and cDNA experiments were quantified using Array Vision software (Imaging Research). Subsequent analysis of the data was performed as described (33). The experiments were always done in triplicate. To calculate synthesis rate (SR) the transcription rates were normalized by the cellular volume (Figure S2C) (58).

More detailed procedures can be found in SI Appendix.

### Chromatin association assay

The chromatin association assay experiment was performed as previously described in (32), with modifications. Yeast cell cultures were grown in 300 mL of YPD medium at 25°C to mid-log phase (OD_600_ 0.5). Cells were collected and pellets were processed to break cells as described in SI Appendix. The lysate was divided into two samples: one half was treated with RNase A and RNase T1; and the other half was incubated without RNase. After 1-hour incubation at 25°C, chromatin was isolated by three-times centrifugation at 13,000 rpm for 20 min. Chromatin was solubilized and analysed by SDS-PAGE and Western blotting against eIF5A, cytoplasmic Pgk1 and nuclear H4 proteins with specific antibodies. At least three biological replicates of each sample were analyzed. More detailed procedures can be found in SI Appendix.

## Data Availability

ChIP-seq data is deposited in NCBI’s Gene Expression Omnibus (GEO) under accession number GSE303994. The GRO data is deposited in GEO under accession number GSE302334.

## Acknowledgements

Grants PID2020-120066RB-I00 and PID2023-152214NB-100 funded by MCIN/AEI/10.13039/501100011033 to PA and PID2020-112853GB-C31 to JEP-O. This research was also funded by Generalitat Valenciana (AICO/2020/086 to PA and CIAICO/2022/237 to PA and JEP-O). MB-A. was a recipient of a predoctoral fellowship (FPU2017/03542) funded by MCIN /AEI/10.13039/501100011033. We thank the Functional Genomics Core Facility of the Institute for Research in Biomedicine (IRB, Barcelona) and the Statistics and data analyses Service of the SCSIE-Universitat de València for the ChIP-seq analysis. Authors acknowledge support by all members of egeDtoP lab.

---

**Figure S1.**
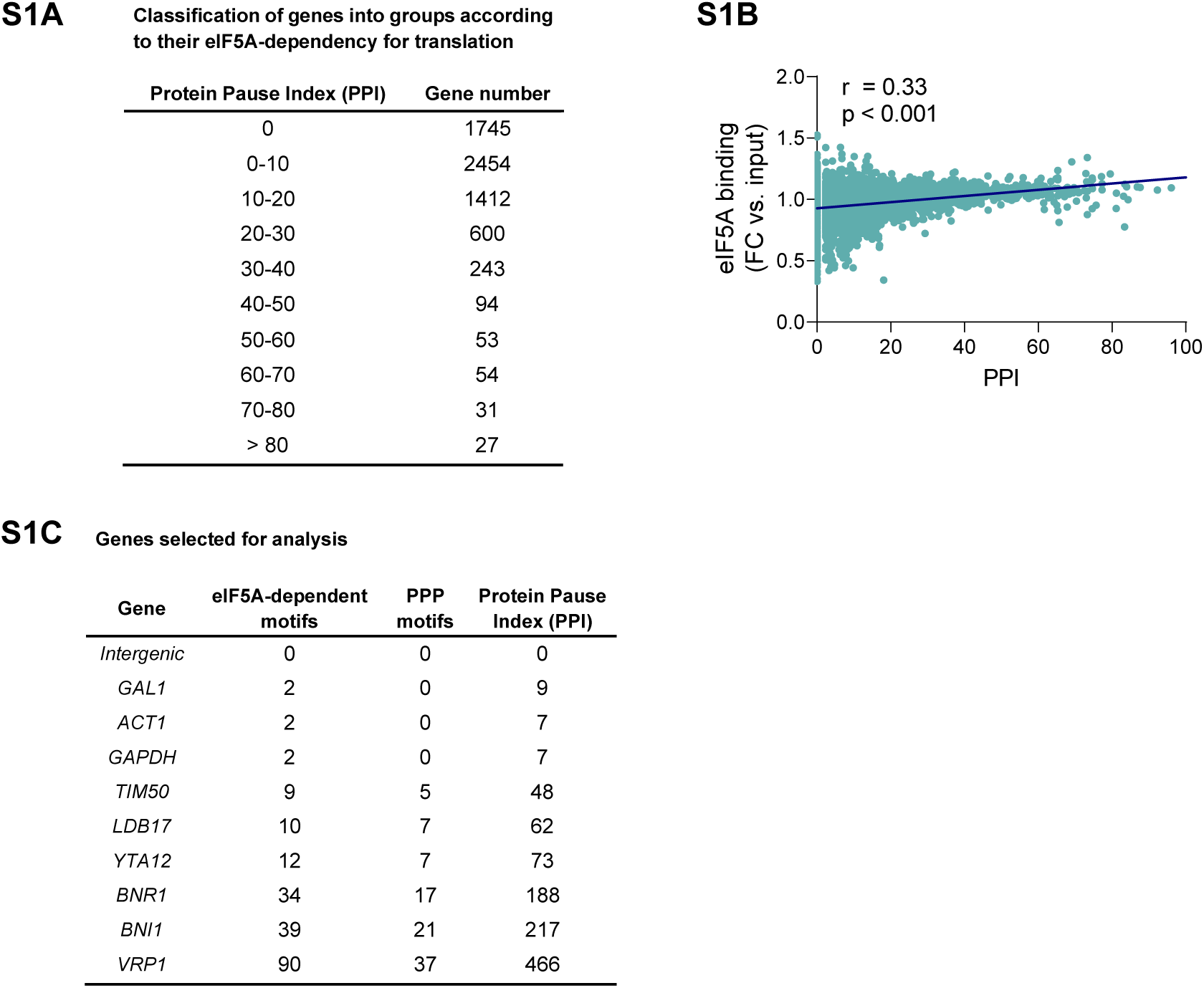
eIF5A chromatin binding positively correlates with the protein pause index (PPI) (A) Classification of genes into groups with respect their putative dependency on eIF5A for their translation as determined by the different PPI intervals (see M&M for full definition of PPI). The number of genes included in each group is shown. (B) eIF5A binding values from ChIP-seq analyses were plotted against the protein pause index (PPI) value associated for each gene. Experimental data were adjusted to potential trend. The Pearson’s correlation coefficient and the associated significance for the plot is shown.(C) Genes selected for ChIP analysis. Number of total eIF5A-dependent tripeptide motifs (Pelechano and Alepuz, 2017), polyproline motifs (PPP) and PPI indexes are shown.

**Figure S2.**
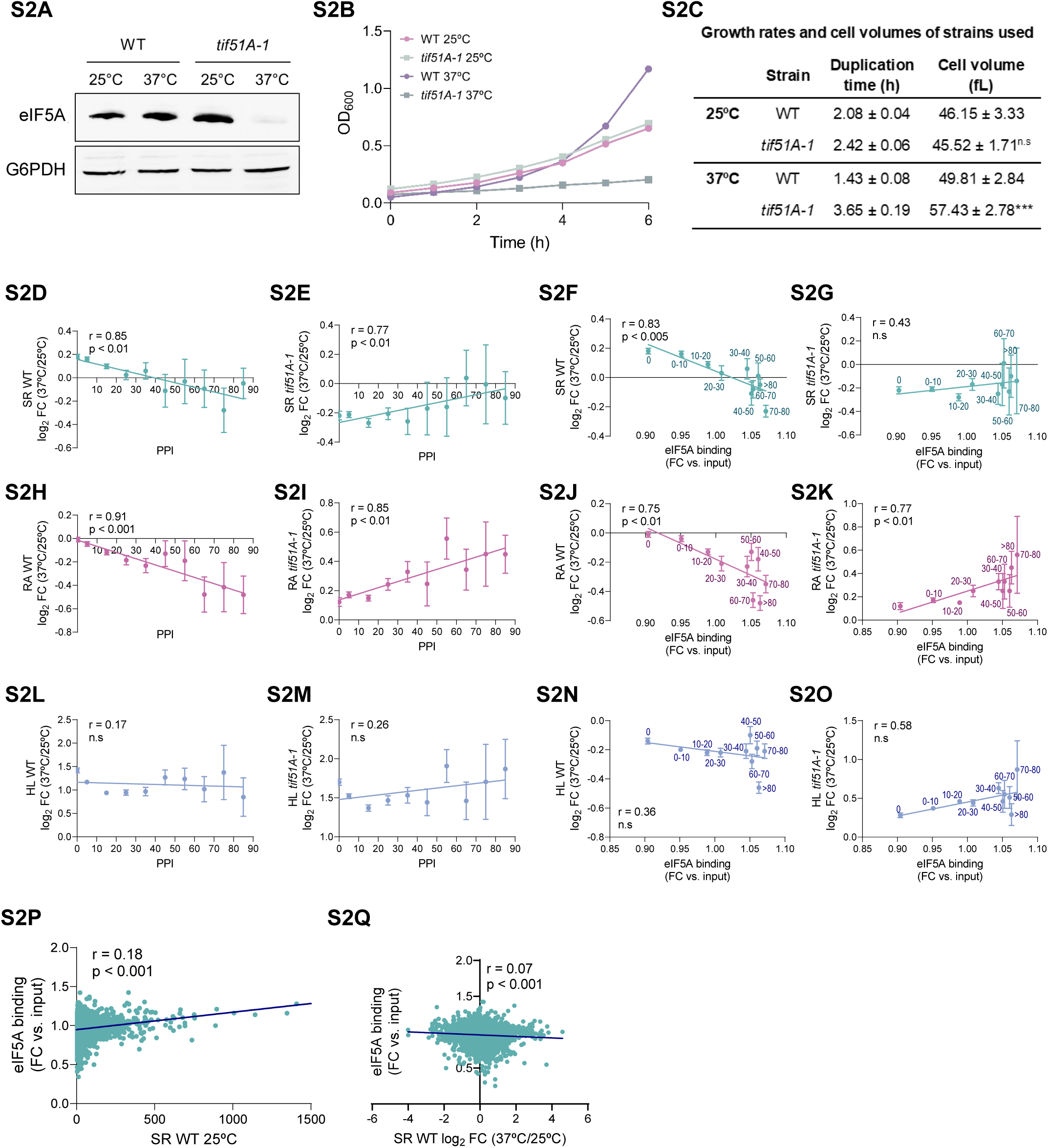
Changes in synthesis rate (SR) and RNA amount (RA) following eIF5A depletion are correlated with the extent to which translation is dependent on eIF5A and with eIF5A binding to chromatin. (A,B,C) eIF5A depletion causes changes in growth rate. Wild-type and *tif51A-1* mutant yeast strains were grown in YPD medium at 25°C until early exponential phase and then transferred to 25°C and 37°C. (A) A representative Western blotting experiment from three independent experiments of the eIF5A protein levels in the wild-type and *tif51A-1* cells incubated at the indicated temperatures for 4 hours. G6PDH protein levels were used as loading control. (B) Growth measured as OD_600_ of the yeast cultures at the indicated time points. A representative experiment from three independent replicates is shown. (C) Duplication times were calculated from data in (B). Cell volume was measured with a Coulter counter for each strain at the corresponding temperature. Statistical significance for volume differences was determined using a two-tailed paired Student’s t-test relative to wild-type cells at 25°C. ***p<0.001. n.s means no significant differences. (D,E,H,I,L,M) Graphs represent the average SR (D,E), RA (H,I), and half-lives (HL) (L,M) for all genes included in each Protein Pause Index (PPI) interval group. Values are given as the log_2_ Fold Change (FC) at 37°C vs. 25°C ± S.E. of wild-type or *tif51A-1* cells. (F,G,J,K,N,O) Graphs represent the average SR (F,G), RA (J,K), and HL (N,O) for all genes included in different PPI group (the labels indicate the PPI interval) versus the average eIF5A binding from ChIP-seq analyses for each group of genes. Values represent the log_2_ Fold Change (FC) at 37°C vs. 25°C ± S.E. of wild-type or *tif51A-1* strain. (P) eIF5A binding values from ChIP-seq analyses were plotted against the corresponding SR value from GRO analysis associated for each gene. (Q) eIF5A binding values from ChIP-seq analyses were plotted against the log_2_ Fold Change at 37°C vs. 25°C of SR of wild-type cells. (D-Q) Experimental data were adjusted to linear trends. Pearson’s correlation coefficient and the associated significance for the plots are shown. n.s means no significant differences.

**Figure S3.**
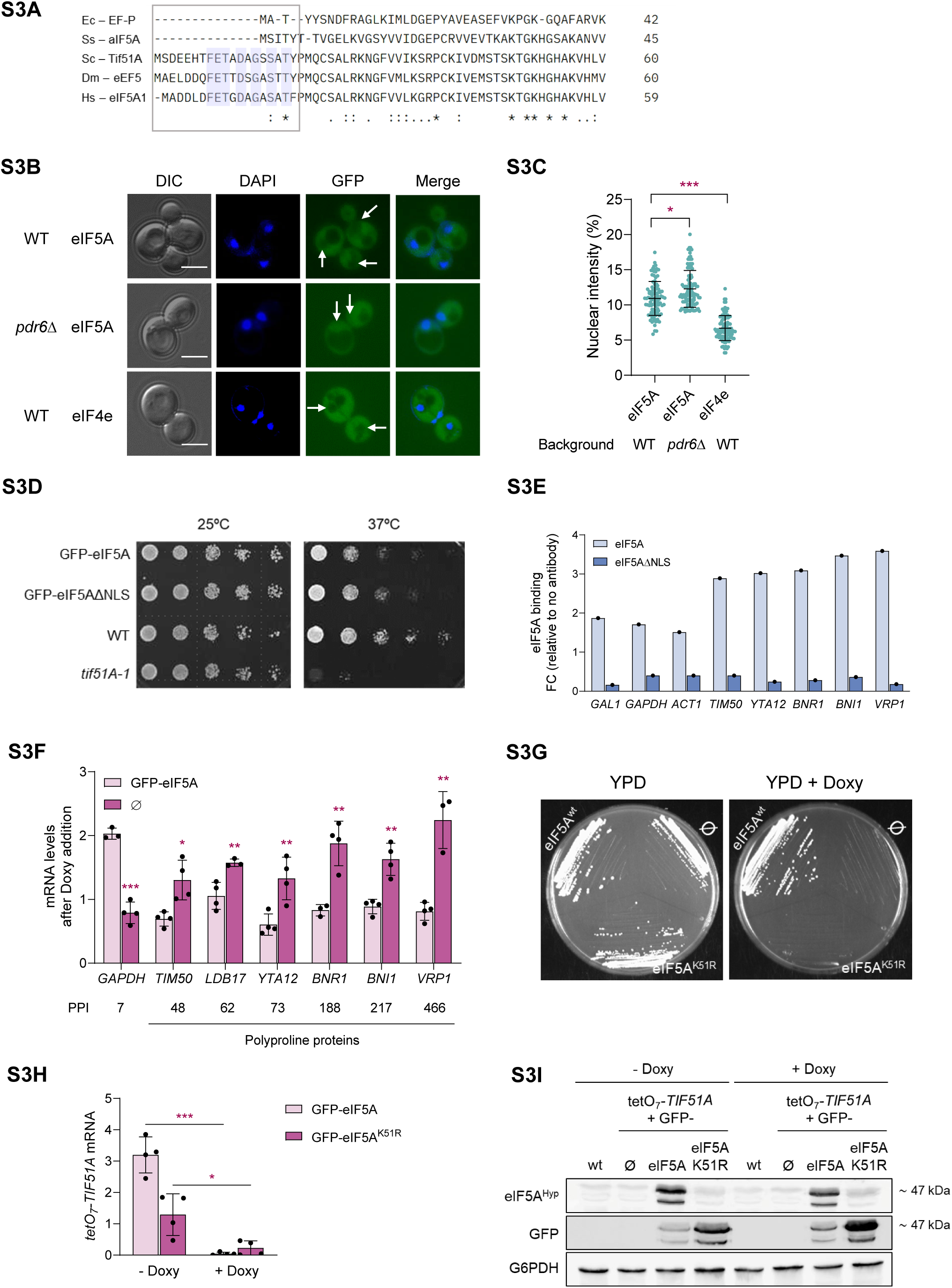
Localization, expression and functionality of yeast strains with modified eIF5A nuclear localization and cytoplasmic activity. (A) Amino acid sequence alignment of genes encoding EF-P in *E.coli* (Ec-EF-P), eIF5A in archaeal *Saccharolobus solfataricus* (S-aIF5A), Tif51A in *S. cerevisiae,* eIF5A in *Drosophila melanogaster* (Dm-eEF5) and eIF5A1 in *Homo sapiens* (H.s). Only the protein N-terminal regions are shown. Residues with an asterisk are conserved in all organisms and residues in purple are identical only in the eukaryotic eIF5A. The region containing the first 19 amino acid residues responsible for the nuclear localization is marked by a box. (B) Wild-type and *pdr6*Δ strains expressing a second copy of GFP-eIF5A, and wild-type strain expressing the control eIF4e-GFP were cultured in YPD medium until reaching exponential phase and subjected to fluorescence microscopy. Cells were incubated for 5 mins with DAPI prior microscopy to stain the nuclei. White arrows indicate the nuclei. A representative image is shown from three independent experiments. Scale bar, 4 μm. (C) Quantification of percentage of nuclear signal is shown from a minimum of 100 cells. Results are presented as individual values together with the mean ± SD from three independent experiments. The statistical significance was measured by using a two-tailed paired Student t-test. (D) Wild-type expressing GFP-eIF5A or GFP-eIF5AΔNLS in the *TIF51A* locus, wild-type and *tif51A-1* strains were cultured in YPD plates at the indicated temperatures. (E) ChIP analysis of eIF5A recruitment in wild-type expressing GFP-eIF5A or GFP-eIF5AΔNLS in the *TIF51A* locus and exponentially-grown in YPD at 25°C. ChIP of eIF5A was performed using an anti-eIF5A antibody. The immunoprecipitated DNA was used to quantify the binding to different genes by qPCR using primers designed for amplification in the ORF regions. The percentage of the signal obtained in each ChIP sample with respect to the signal obtained with the DNA from the corresponding whole cell extract was calculated. One representative experiment is shown. (F) Yeast strains with tetO_7_-*TIF51A* at the *TIF51A* locus and with a second copy of GFP-eIF5A or without second copy (Ø) were cultured overnight in YPD at 25°C in the presence of doxycycline (15 µM) to deplete eIF5A expressed from the first copy, and then, mRNA levels from each gene were determined by RT-qPCR using primers designed for amplification in the ORF regions. (G) Yeast strain with tetO_7_-*TIF51A* at the endogenous locus and with a second copy of GFP-eIF5A, GFP-eIF5A^K51R^ or without second copy (Ø) were grown in YPD and then plated to test growth in YPD plates with or without doxycycline (15 µM). (H,I) Same yeast strains used in (G) were cultured overnight in YPD at 25°C in the presence or absence of doxycycline (15 µM) and then, the tetO_7_-*TIF51A* mRNA levels were determined by RT-qPCR using specific primers (H), and the expression of the second copy GFP-eIF5A, of GFP-eIF5A*^K51R^* was tested by Western blotting using anti-hypusinated-eIF5A, anti-GFP or anti-G6PDH antibodies. A wild-type strain was included as additional negative control (I). (F,H) Results are presented as individual values together with the mean ± SD from at least three independent experiments. Statistical significance was determined using a two-tailed paired Student’s t-test. *p<0.05, **p<0.01, ***p<0.001. n.s means no significant differences.

**Figure S4.**
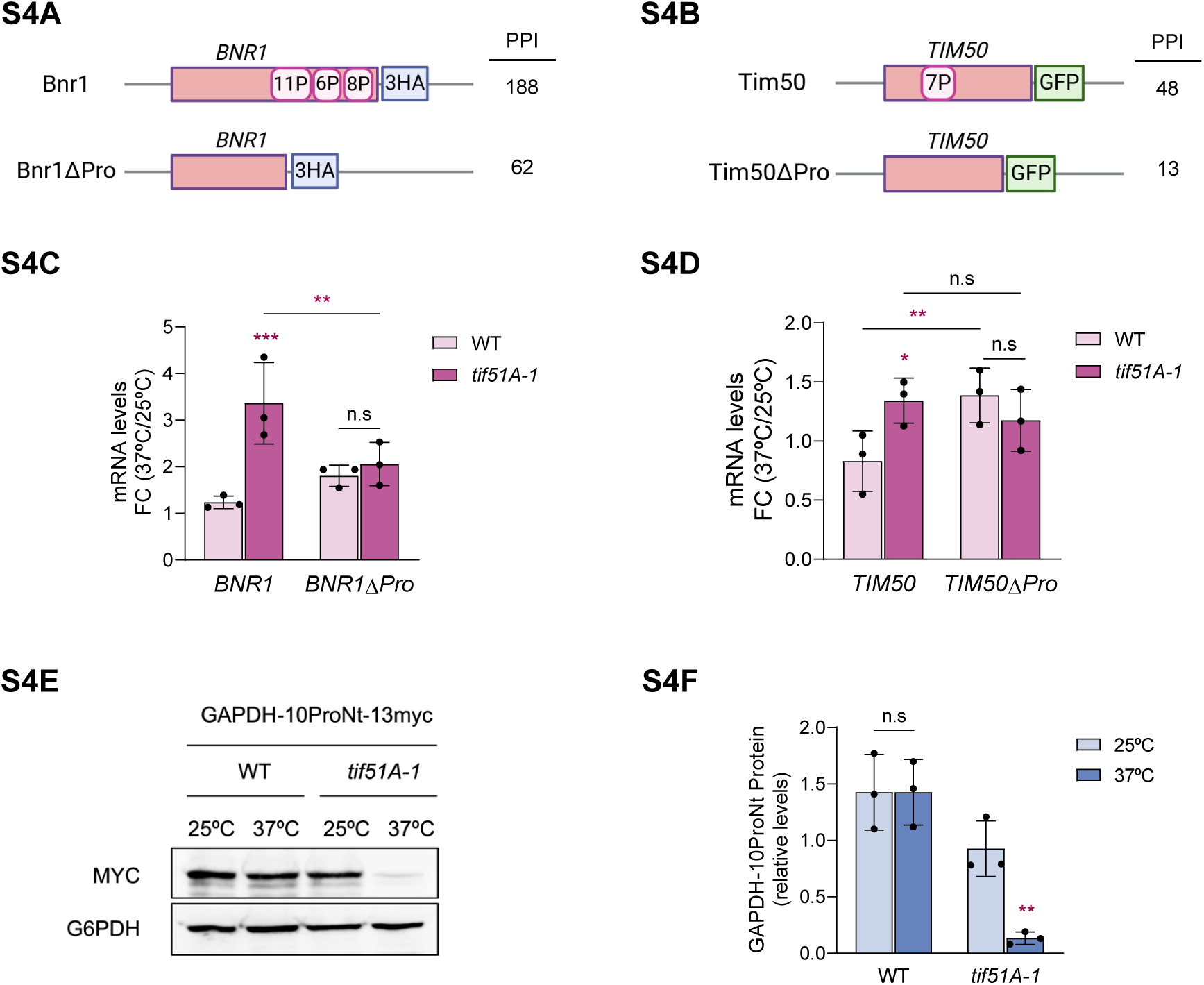
Nucleotide sequences that encode polyPro motifs are necessary to promote transcriptional repression by eIF5A. (A,B) Schematic diagram showing the C-terminal genomic tagging of native *BNR1* (A) and *TIM50* (B) ORFs as well as their polyPro-deleted versions (*BNR1ΔPro, TIM50ΔPro*). PPI of both native and mutated versions are shown. (C,D) Exponentially growing cultures of wild-type and *tif51A-1* mutant yeast strains, carrying native and mutated versions of *BNR1* and *TIM50* genes were grown in YPD medium at 25°C until exponential phase and then transferred to 25°C and 37°C for four hours. mRNA levels of the *BNR1* (C) and *TIM50* (D) versions were determined by RT-qPCR. (C,D) Data are presented as the mean fold change (FC) 37°C vs. 25°C ± SD from three independent experiments. Statistical significance was determined using a two-tailed paired Student’s t-test relative to corresponding wild-type cells. (E,F) Wild-type and *tif51A-1* yeast strains carrying a genomic *GAPDH* version with an N-terminal insertion of a polyPro sequence and a C-terminal tagging with myc (*GAPDH*-10ProNt-13myc, see schematic diagram in Fig.4D) were grown in YPD medium at 25°C until exponential phase and then transferred to 25°C and 37°C for four hours. Protein levels of the Gapdh protein with the polyPro sequence were determined by Western blotting using anti-myc antibodies and anti-G6PDH as loading control (E), and quantified (F). (F) Data are shown as the mean relative protein level ± SD from three independent experiments. Statistical significance of protein level at 37°C vs. 25°C was determined using a two-tailed paired Student’s t-test. *p<0.05, **p<0.01, ***p<0.001. n.s means no significant differences.

**Figure S5.**
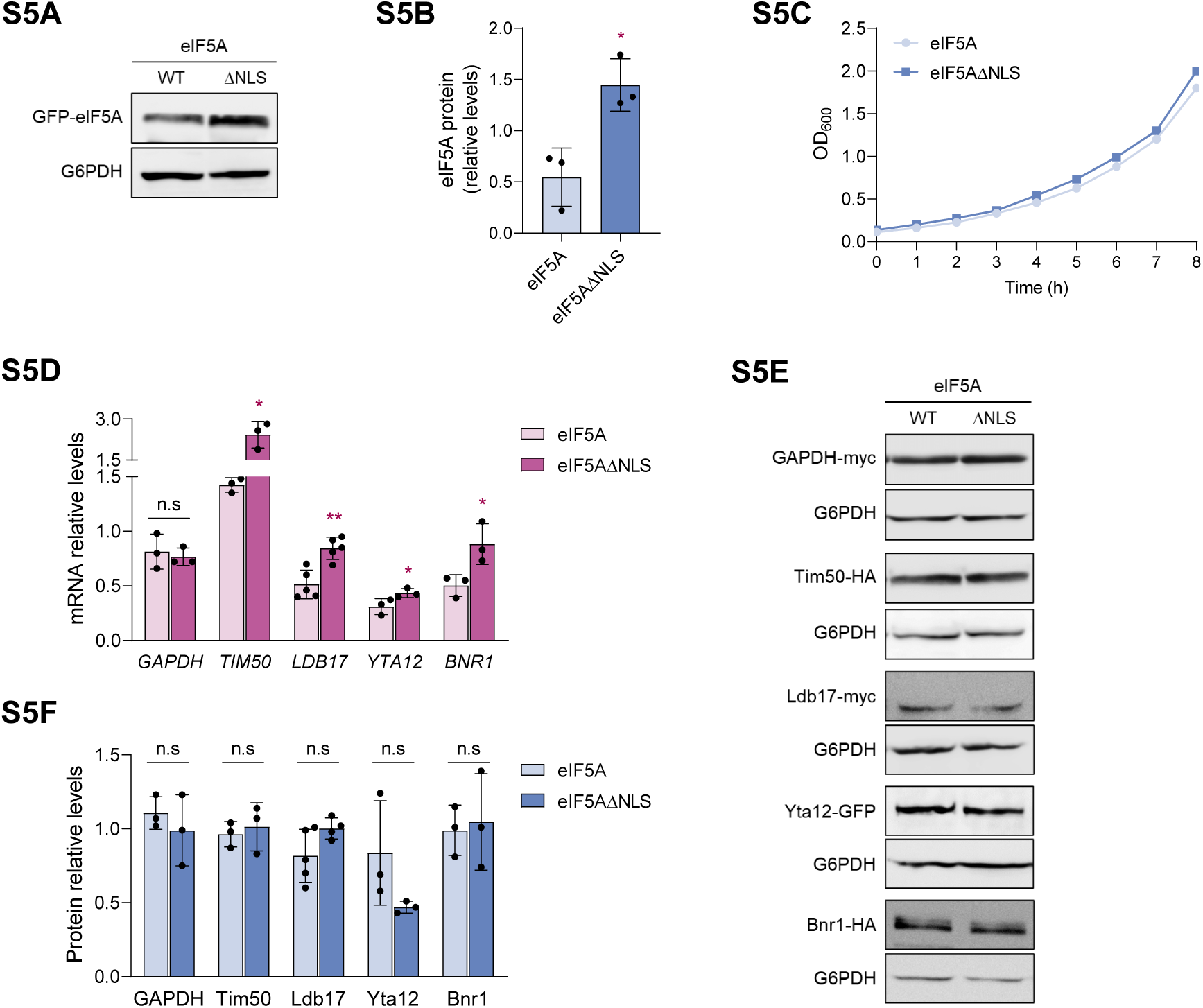
Study of the mRNA and protein synthesis by an eIF5A mutant protein without the NLS region. (A-F) Wild-type cells expressing the native eIF5A or the eIF5AΔNLS version in the *TIF51A* locus were grown in YPD medium at 30°C until exponential phase. eIF5A protein levels were determined by Western blotting (A) and quantified from three independent replicates (B) and growth was determined by measuring the OD_600_ of the yeast cultures at the indicated time points. A representative experiment is shown (C). (D) mRNA levels from *GAPDH*, *TIM50*, *LDB17*, *YTA12* and *BNR1* genes were determined by RT-qPCR using primers designed for amplification in the specific ORF regions (E,F) The corresponding protein levels were determined by Western blotting (E) and quantified (F). (A,E) G6PDH levels were used as loading control. A representative image is shown. (B,D,F) Data are presented as the mean ± SD from at least three independent experiments. Statistical significance was determined using a two-tailed paired Student’s t-test relative to corresponding wild-type cells. *p<0.05, **p<0.01. n.s means no significant differences.

## Notes

### Competing Interest Statement

The authors have declared no competing interest.

