## Supplementary Tables for "eIF5A coordinates the transcription and translation of its target genes"

**Table S1. Yeast strains used in this study**

| Name | Genotype | Source |
|---|---|---|
| **BY4741** | MATa *ura3Δ0 leu2Δ0 his3Δ1 met15Δ0* | Euroscarf |
| ***tif51A-1*** | BY4741 MATa *ura3Δ0 leu2Δ0 his3Δ1 met15Δ0 tif51A-1::kanR* | (Li *et al.,* 2011) |
| **PAY666** | BY4741 MATa *ura3Δ0 leu2Δ0 his3Δ1 met15Δ0 eIF4e-GFP-his3MX6* | This study |
| **PAY778** | BY4741 MATa ura3Δ0 leu2Δ0 his3Δ1 met15Δ0 $tetO_7$*-TIF51A-kanMX* | This study |
| **PAY888** | BY4741 MATa *ura3Δ0 leu2Δ0 his3Δ1 met15Δ0 TIM50-GFP-his3MX6* | (Barba-Aliaga *et al.*, 2024) |
| **PAY890** | BY4741 MATa *ura3Δ0 leu2Δ0 his3Δ1 met15Δ0 tif51A-1::kanR TIM50-GFP-his3MX6* | (Barba-Aliaga *et al.*, 2024) |
| **PAY892** | BY4741 MATa *ura3Δ0 leu2Δ0 his3Δ1 met15Δ0 YTA12-GFP-his3MX6* | (Barba-Aliaga *et al.*, 2024) |
| **PAY893** | BY4741 MATa *ura3Δ0 leu2Δ0 his3Δ1 met15Δ0 tif51A-1::kanR YTA12-GFP-his3MX6* | (Barba-Aliaga *et al.*, 2024) |
| **PAY896** | BY4741 MATa *ura3Δ0 leu2Δ0 his3Δ1 met15Δ0 BNR1-3HA-his3MX6* | This study |
| **PAY898** | BY4741 MATa *ura3Δ0 leu2Δ0 his3Δ1 met15Δ0 tif51A-1::kanR BNR1-3HA-his3MX6* | This study |
| **PAY899** | BY4741 MATa *ura3Δ0 leu2Δ0 his3Δ1 met15Δ0 BNR1ΔPro-3HA-his3MX6* | This study |
| **PAY901** | BY4741 MATa *ura3Δ0 leu2Δ0 his3Δ1 met15Δ0 tif51A-1::kanR BNR1ΔPro-3HA-his3MX6* | This study |
| **PAY1083** | BY4741 MATa *ura3Δ0 leu2Δ0 his3Δ1 met15Δ0 GFP-TIF51A-his3MX6* | This study |
| **PAY1085** | BY4741 MATa *ura3Δ0 leu2Δ0 his3Δ1 met15Δ0 TIM50ΔPro-GFP-his3MX6* | This study |
| **PAY1086** | BY4741 MATa *ura3Δ0 leu2Δ0 his3Δ1 met15Δ0 tif51A-1::kanR TIM50ΔPro-GFP-his3MX6* | This study |
| **PAY1087** | BY4741 MATa *ura3Δ0 leu2Δ0 his3Δ1 met15Δ0 YTA12ΔPro-GFP-his3MX6* | This study |
| **PAY1088** | BY4741 MATa *ura3Δ0 leu2Δ0 his3Δ1 met15Δ0 tif51A-1::kanR YTA12ΔPro-GFP-his3MX6* | This study |
| **PAY1443** | BY4741 MATa *ura3Δ0 leu2Δ0 his3Δ1 met15Δ0 GAPDH-13myc-his3MX6* | This study |
| **PAY1444** | BY4741 MATa *ura3Δ0 leu2Δ0 his3Δ1 met15Δ0 tif51A-1::kanR GAPDH-13myc-his3MX6* | This study |
| **PAY1631** | BY4741 MATa *ura3Δ0 leu2Δ0 his3Δ1 met15Δ0 GFP-TIF51A*$^{K51R}$*-his3MX6* | This study |
| **PAY1632** | BY4741 MATa *ura3Δ0 leu2Δ0 his3Δ1 met15Δ0 GFP-TIF51AΔNLS-his3MX6* | This study |
| **PAY1633** | BY4741 MATa *ura3Δ0 leu2Δ0 his3Δ1 met15Δ0 pdr6::kanMX6 GFP-TIF51A-his3MX6* | This study |
| **PAY1673** | BY4741 MATa *ura3Δ0 leu2Δ0 his3Δ1 met15Δ0 tif51A-1::kanR GFP-TIF51A-his3MX6* | This study |
| **PAY1675** | BY4741 MATa *ura3Δ0 leu2Δ0 his3Δ1 met15Δ0 tif51A-1::kanR GFP-TIF51A*$^{K51R}$*-his3MX6* | This study |
| **PAY1678** | BY4741 MATa *ura3Δ0 leu2Δ0 his3Δ1 met15Δ0 tif51A-1::kanR GFP-TIF51AΔNLS-his3MX6* | This study |
| **PAY1759** | BY4741 MATa *ura3Δ0 leu2Δ0 his3Δ1 met15Δ0 tif51A::GFP-TIF51A-his3MX6* | This study |
| **PAY1760** | BY4741 MATa *ura3Δ0 leu2Δ0 his3Δ1 met15Δ0 tif51A::GFP-TIF51AΔNLS-his3MX6* | This study |
| **PAY1915** | BY4741 MATa *ura3Δ0 leu2Δ0 his3Δ1 met15Δ0 tif51A::GFP-TIF51A-his3MX6 LDB17-13myc-KanMX* | This study |
| **PAY1916** | BY4741 MATa *ura3Δ0 leu2Δ0 his3Δ1 met15Δ0 tif51A::GFP-TIF51AΔNLS-his3MX6 LDB17-13myc-KanMX* | This study |
| **PAY1917** | BY4741 MATa *ura3Δ0 leu2Δ0 his3Δ1 met15Δ0 tif51A::GFP-TIF51A-his3MX6 BNR1-3HA-KanMX* | This study |
| **PAY1919** | BY4741 MATa *ura3Δ0 leu2Δ0 his3Δ1 met15Δ0 tif51A::GFP-TIF51AΔNLS-his3MX6 BNR1-3HA-KanMX* | This study |
| **PAY1927** | BY4741 MATa *ura3Δ0 leu2Δ0 his3Δ1 met15Δ0 tif51A::GFP-TIF51A-his3MX6 GAPDH-13myc-KanMX* | This study |
| **PAY1928** | BY4741 MATa *ura3Δ0 leu2Δ0 his3Δ1 met15Δ0 tif51A::GFP-TIF51AΔNLS-his3MX6 GAPDH-13myc-KanMX* | This study |
| **PAY1948** | BY4741 MATa *ura3Δ0 leu2Δ0 his3Δ1 met15Δ0 tif51A::GFP-TIF51A-his3MX6 YTA12-GFP-KanMX* | This study |
| **PAY1949** | BY4741 MATa *ura3Δ0 leu2Δ0 his3Δ1 met15Δ0 tif51A::GFP-TIF51AΔNLS-his3MX6 YTA12-GFP-KanMX* | This study |
| **PAY1950** | BY4741 MATa *ura3Δ0 leu2Δ0 his3Δ1 met15Δ0 tif51A::GFP-TIF51A-his3MX6 TIM50-3HA-KanMX* | This study |
| **PAY1952** | BY4741 MATa *ura3Δ0 leu2Δ0 his3Δ1 met15Δ0 tif51A::GFP-TIF51AΔNLS-his3MX6 TIM50-3HA-KanMX* | This study |

| | | |
|---|---|---|
| **PAY2118** | BY4741 MATa *ura3Δ0 leu2Δ0 his3Δ1 met15Δ0 tetO*$_7$*-TIF51A-kanMX GFP-TIF51A-his3MX6* | This study |
| **PAY2130** | BY4741 MATa *ura3Δ0 leu2Δ0 his3Δ1 met15Δ0 GAPDH-10ProNt-13myc-his3MX6* | This study |
| **PAY2132** | BY4741 MATa *ura3Δ0 leu2Δ0 his3Δ1 met15Δ0 tif51A-1::kanR GAPDH-10ProNt-13myc-his3MX6* | This study |
| **PAY2158** | BY4741 MATa *ura3Δ0 leu2Δ0 his3Δ1 met15Δ0 tetO*$_7$*-TIF51A-kanMX GFP-TIF51A*$^{K51R}$*-his3MX6* | This study |
| **PAY2246** | BY4741 MATa *ura3Δ0 leu2Δ0 his3Δ1 met15Δ0 CTK1-GFP-his3MX6* | This study |

**Table S2. Plasmids used in this study**

| Name | Plasmid description | Source |
|---|---|---|
| **PA201** | pFA6a-3HA-HIS3MX6 | (Longtine *et al.,* 1998) |
| **PA234** | pFA6a-13myc-HIS3MX6 | (Longtine *et al.,* 1998) |
| **PA235** | pFA6a-GFP-kanMX6 | (Longtine *et al.,* 1998) |
| **PA239** | pFA6a-13myc-kanMX6 | (Longtine *et al.,* 1998) |
| **PA241** | pFA6a-3HA-kanMX6 | (Longtine *et al.,* 1998) |
| **PA242** | pFA6a-GFP-HIS3MX6 | (Longtine *et al.,* 1998) |
| **PA410** | pRS303-peIF5A-GFP-TIF51A-HIS3MX6 | Dr. Brian M. Zid |
| **PA431** | pRS303-peIF5A-GFP-TIF51A$^{K51R}$-HIS3MX6 | This study |
| **PA432** | pRS303-peIF5A-GFP-eIF5AΔNLS-HIS3MX6 | This study |

**Table S3. Oligonucleotides used in this study**

| Primer | Sequence (5'-3') | |
|---|---|---|
| **Gene expression detection by RT-qPCR and ChIP-qPCR** | | |
| ACT1-F | TCGTTCCAATTTACGCTGGTT | RT-qPCR/ChIP-qPCR |
| ACT1-R | CGGCCAAATCGATTCTCAA | RT-qPCR/ChIP-qPCR |
| BNI1-F | ACATGTGGAAAACGGAAAGC | RT-qPCR/ChIP-qPCR |
| BNI1-R | AGATCTTCTGCGCCATCTGT | RT-qPCR/ChIP-qPCR |
| BNR1-F | CCAGCTCCACCTTTACCAAA | RT-qPCR/ChIP-qPCR |
| BNR1-R | CCCAGTGGATTTGCTTCAAT | RT-qPCR/ChIP-qPCR |
| GAL1-F | TGGTGTTAACAATGGCGGTA | ChIP-qPCR |
| GAL1-R | GGGCGGTTTCAAACTTGTTA | ChIP-qPCR |
| GAPDH-F | ATGACCGCCACTCAAAAGAC | RT-qPCR/ChIP-qPCR |
| GAPDH-R | CTTAGCAGCACCGGTAGAGG | RT-qPCR/ChIP-qPCR |
| Intergenic-F | GGCTGTCAGAATATGGGGCCGTAGTA | ChIP-qPCR |
| Intergenic-R | CACCCCGAAGCTGCTTTCACAATAC | ChIP-qPCR |
| LDB17-F | AACTCAAGCCTGGTTGCCTA | RT-qPCR/ChIP-qPCR |
| LDB17-R | CATCCGGTAGAGGTCACGAT | RT-qPCR/ChIP-qPCR |
| tetO-TIF51 qPCR | AATTACCGGATCAATTCGGG | RT-qPCR |
| TIF51A-7-R | GTGTATGCTTTAGGAGAACG | RT-qPCR |
| TIM50-F | TCTGCGTTGACAGGTACTGC | RT-qPCR/ChIP-qPCR |
| TIM50-R | AATCAGGGAAAGGTGGCTCT | RT-qPCR/ChIP-qPCR |
| VRP1-F | GGCAGAAATTAATGCCAGGA | RT-qPCR/ChIP-qPCR |
| VRP1-R | GTGGTGCAGTAGGCGGTAAT | RT-qPCR/ChIP-qPCR |
| YTA12-F | GTTTGTTGGTGTTGGTGCAG | RT-qPCR/ChIP-qPCR |
| YTA12-R | TCATCATTGGCACCTGAAAA | RT-qPCR/ChIP-qPCR |
| **Gene tagging by PCR** | | |
| BNR1-F2 | TTACTAGAGAGAACGCATGCTATGCTGAACGATATTCAAAATATACGGATCCCCGGGTTAATTAA | |
| BNR1-R1 | TTTCTTTATATAAGCTCCACAACTACATAAAATACTAAGTCTTCAGAATTCGAGCTCGTTTAAAC | |
| CTK1-Tagging-F | AATAGTAATAATAATAATAATAATAATAATGACGATGATGATAAACGGATCCCCGGGTTAATTAA | |
| CTK1-Tagging-R | TTAATCTATTTTTTGTGTCTACTTATTTCAATTGGCTATATATCCGAATTCGAGCTCGTTTAAAC | |
| GAPDH-F2 | TACTCCGCCAGAGTTGTTGACTTGATCGAATATGTTGCCAAGGCTCGGATCCCCGGGTTAATTAA | |
| GAPDH-R1 | TGTATATTCAAAAAAAAATCATTATCCTCATCAAGATTGCTTTATGAATTCGAGCTCGTTTAAAC | |
| GFP-eIF5A-F | TAGACTCCCAAACACACACAAATACCAACTCATATATACACTAGTACACTCTATATTTTTTTATG | |
| GFP-eIF5A-R | TCTTTTTTCATTTATATCCCATGCCATGATGTTAACCGGTTTAATCGGTTCTAGCAGCTTCCTTG | |
| LDB17-F2 | CCGCCTCCTCCTCCTCCTCCATCAAGAAAATGTGGAACTCCAAAACGGATCCCCGGGTTAATTAA | |
| LDB17-R1 | ATGGTCGGAAGAACCACATTAATGCAAGAAGAAATAATGCTTTACGAATTCGAGCTCGTTTAAAC | |
| TIM50-F2 | TTATTTGAAGAGGAAAAGAAAAAAGAAGAAGATTGCTGAATCCAAACGGATCCCCGGGTTAATTAA | |
| TIM50-R1 | CACACATAGATACGTAGATACATGAGAAGAGGGTTTACATGAAAAGAATTCGAGCTCGTTTAAAC | |
| YTA12-F2 | GAAGAAAAAAACGAAAAACGTAATGAGCCTAAGCCATCTACAAACCGGATCCCCGGGTTAATTAA | |
| YTA12-R1 | ATATGTAGAACAGTCTTCCTCCATTTCTTTGTATTGTGAAATATCGAATTCGAGCTCGTTTAAAC | |

| | |
|---|---|
| **Proline deletion/insertion by PCR** | |
| BNR1-delPro-F | AGTCTTGATAATGGAATCCAACTAGTACCTGAAGTTGTTAAACTATCCTTGTCGATGAACAAA |
| BNR1-delPro-R | TTTCTTTATATAAGCTCCACAACTACATAAAATACTAAGTCTTCAGAATTCGAGCTCGTTTAAAC |
| TIM50-delPro-F | CCTACTTCCAAGAGCCACCTTTCCCTGATTTACTACCAAAGGCCATTAACTCTTG |
| TIM50-delPro-R | CACACATAGATACGTAGATACATGAGAAGAGGGTTTACATGAAAA |
| YTA12-delPro-F | GCTACTTTGAAGGTAACAATAGCAGAAATATTCCACTAAATGATCCTAGTAATCC |
| YTA12-delPro-R | TTTCTTTATATAAGCTCCACAACTACATAAAATACTAAGTCTTCA |
| GAPDH-10ProNt-F3 | CCACCGCCTCCTCCTCCTCCTCCTCCACCCTTCGGTAGAATCGGTAGATT |
| GAPDH-10ProNt-F4 | CACTAAATTTACACACAAAACAAAATGATCAGAATTGCTATTAACGGTCCACCGCCTCCTCCTCCTCC |
| **Targeted mutagenesis of PA410 plasmid** | |
| K51R-F | TAAGACTGGTCGCCACGGTCACG |
| K51R-R | GAAGTGGACATGTCG |
| ΔNLS-F | GGATGAACTATACAACCAATGCAATGTTCTGCCTTGAGAA |
| ΔNLS-R | TTGTATAGTTCATCCATGCCATGTG |

**Table S4. Antibodies used in this study**

| Primary Antibody | Working dilution | Source | Secondary Antibody | Working dilution | Source |
|---|---|---|---|---|---|
| **Protein detection by Western blotting** | | | | | |
| eIF5A | 1:500 | Abcam (Ab32407) | α-rabbit | 1:10000 | Promega |
| G6PDH | 1:15000 | Roche (10127671001) | α-rabbit | 1:10000 | Promega |
| HA | 1:5000 | Roche (12013819001) | - | - | - |
| hyp-eIF5A | 1:600 | Genentech (FabHpu) | α-rabbit | 1:10000 | Promega |
| Myc | 1:1000 | Invitrogen (13-2500) | α-mouse | 1:10000 | Promega |
| GFP | 1:5000 | Merck (11814460001) | α-mouse | 1:10000 | Promega |
| H4 | 1:1000 | Abcam (Ab7311) | α-rabbit | 1:10000 | Promega |
| PGK1 | 1:10000 | ThermoFisher (22C5D8) | α-mouse | 1:10000 | Promega |
| **Chromatin immunoprecipitation** | | | | | |
| eIF5A | 1:4 | Abcam (Ab32407) | Dynabeads anti-rabbit IgG | | Invitrogen (11203D) |
| Rpb1 (8WG16) | 1:7 | Invitrogen (MA1-10882) | Dynabeads Pan Mouse IgG | | Invitrogen (11041) |
