## Supplementary M&M for "eIF5A coordinates the transcription and translation of its target genes"

### Supplementary Information Appendix 1.

#### Materials and methods

#### Yeast strains, plasmids, and growth conditions

All *Saccharomyces cerevisiae* strains and plasmids used herein are listed in Supplementary Tables S1 and S2 respectively. For all the experiments carried out *S. cerevisiae* cells were grown in liquid YPD (2% glucose, 2% peptone, 1% yeast extract).

Plasmid pRS303-peIF5A-GFP-TIF51A-HIS3MX6 (PA410) and derivatives were used for expression of a second copy of eIF5A fused to GFP and subsequent visualization under microscopy. Plasmids pRS303-peIF5A-GFP-TIF51A$^{K51R}$-HIS3 (PA431) and pRS303-peIF5A-GFP-TIF51AΔNLS-HIS3MX6 (PA432) were constructed from the plasmid PA410, which was used as a template for PCR reaction using primers listed in Table S3 ("targeted mutagenesis" subsection). Parental plasmid was digested by restriction enzyme DpnI (Thermo Fisher Scientific) and the rest was transformed into bacteria for plasmid amplification and sequencing. The resulting plasmids were integrated into the genome of the corresponding strains by homologous recombination following the lithium acetate-based method (Gietz et al., 1992) and transformants were selected in SC medium lacking histidine.

A PCR-based genomic tagging technique was employed to substitute the genomic full length eIF5A (*TIF51A*) ORF by eIF5A or eIF5AΔNLS fused to GFP at the N-terminal region. The plasmids PA410 and PA432 were used as templates for PCR reaction using primers listed in Table S3 ("gene tagging by PCR" subsection). The resulting cassettes were transformed in the corresponding strains following the lithium acetate-based method (Gietz et al., 1992) and transformants were selected in SC medium lacking histidine. All the integrations were confirmed by genomic DNA conventional PCR.

A PCR-based genomic tagging technique was employed to tag the genomic full length *YTA12, TIM50, BNR1, LDB17* and *GAPDH* (*TDH1*) ORFs with GFP, HA or myc at the C-terminal region. The plasmids pFA6a-GFP-HIS3MX, pFA6a-GFP-KanMX, pFA6a-3HA-HIS3MX, pFA6a-3HA-KanMX, pFA6a-13myc-HIS3MX and pFA6a-13myc-KanMX (Longtine et al., 1998) were used as a template for PCR reaction using primers listed in Table S3 ("gene tagging by PCR" subsection). The resulting cassettes were transformed in the corresponding strains following the lithium acetate-based method (Gietz et al., 1992) and transformants were selected in SC medium lacking histidine or YPD supplemented with geneticin. All the integrations were confirmed by genomic DNA conventional PCR.

To generate the strains harboring the deletion of the consecutive prolines from the *YTA12*, *TIM50* and *BNR1* gene sequences, the C-terminal *YTA12-GFP*, *TIM50-GFP* and *BNR1-3HA* sequences were amplified from genomic DNA of the strain PAY892, PAY888 and PAY896, respectively using primers listed in Table S3 ("proline deletion/insertion by PCR" subsection). The use of these primers resulted in the deletion of nucleotides 463-489 in *YTA12*, 541-561 in *TIM50* and 2305-2478 in *BNR1*, which encode for the 9, 7 and 25 prolines stretch of the Yta12, Tim50 and Bnr1 proteins respectively. The resulting PCR products were transformed in wild-type and *tif51A-1* strains as previously described and transformants were selected in SC medium lacking histidine. To generate the strains harboring the insertion of 10 consecutive prolines at the N-terminal of the *GAPDH* (*TDH1*) gene sequence, plasmid pFA6a-13myc-HIS3MX was used as a template for PCR reaction using primers GAPDH-10ProNt-F3 and GAPDH-10ProNt-F4 (Table S3 "proline deletion/insertion by PCR" subsection). The resulting PCR product was used as a template for a second PCR reaction using primers GAPDH-15Pro-F2 and GAPDH-15Pro-R1 (Table S3 "proline deletion/insertion by PCR" subsection). The second PCR product was then transformed in wild-type and *tif51A-1* strains as previously described and transformants were selected in SC medium lacking histidine. All the insertions and deletions were confirmed by genomic DNA conventional PCR.

Experimental assays were performed with cells exponentially grown for at least four generations until required $OD_{600}$ at the corresponding temperature. Temperature-sensitive strains were grown at the permissive temperature of 25°C until required $OD_{600}$ and transferred to the non-permissive temperature of 37°C for 4 h for complete depletion of eIF5A but maintaining cell viability (Li et al., Genetics 2014).

### RT-qPCR analysis

For the analysis of the mRNA levels, total RNAs were isolated from yeast cells following the phenol:chloroform protocol. Briefly, a volume of an exponential phase culture corresponding to 10 $OD_{600}$ units was harvested and flash frozen. Cells were resuspended in 500 µL of cold LETS buffer (LiCl 0.1 M, EDTA pH 8.0 10 mM, Tris-HCl pH 7.4 10 mM, SDS 0.2%) and transferred into a screw-cap tube already containing 500 µL of sterile glass beads and 500 µL of phenol:chloroform (5:1). Then, cells were broken using the Precellys 24 tissue homogenizer (Bertin Technologies) and centrifuged. The supernatant was transferred into a new tube containing 500 µL of phenol:chloroform (5:1) and then to a tube containing 500 µL of chloroform:isoamyl alcohol (25:1). RNA from the top phase was precipitated and finally dissolved in water for later quantification and quality control with Nanodrop device (Thermo Fisher Scientific).

The reverse transcription and quantitative PCR reactions were performed as detailed in Garre et al., 2013. Briefly, 2.5 µg of the total DNAse-I (Roche) treated RNA were retrotranscribed using an oligo d(T)18 with Maxima Reverse Transcriptase (Thermo Fisher Scientific). cDNA was labelled with SYBR Pre-mix Ex Taq (Tli RNase H Plus, Takara) and the Cq values were obtained from the CFX96 TouchTM Real-Time PCR Detection System (BioRad). Endogenous *ACT1* mRNA levels were used for normalization. At least three biological replicates of each sample were analyzed, and the specific primers designed to amplify gene fragments of interest are listed in Table S3.

### Western blotting

For yeast protein content analysis by western blotting we followed the protocol described in (Zuzuarregui et al., 2015). Briefly, a cell culture volume corresponding to 10 $OD_{600}$ units was harvested by centrifugation. For protein extraction, cell pellets were washed and resuspended in 200 µL of NaOH 0.2M and incubated at room temperature for 5 min for subsequent centrifugation at 12000 rpm for 1 min. Samples were then resuspended in 100 µL of 2X-SDS protein loading buffer (24 mM Tris-HCl pH 6.8, 10% glycerol, 0.8% SDS, 5.76 mM β-mercaptoethanol, 0.04% bromophenol blue) and boiled at 95°C for 5 min. After, lysates were centrifuged at 3000 rpm for 10 min at 4°C to remove cell debris and insoluble proteins, and supernatants were transferred into new tubes and stored at -20°C. Total protein content in the extract was quantified by an $OD_{280}$ estimation in a Nanodrop device (Thermo Fisher Scientific) to load equal protein amounts per sample into the SDS-PAGE gel. The acrylamide percentage of the used SDS-PAGE depended on the molecular weight of the protein of interest.

SDS-PAGE and Western blotting were performed using standard procedures (BioRad). Blotting membranes were blocked with 5% skimmed milk in TBS-T (150 mM NaCl, 20 mM Tris, 0.1% Tween20, pH 7.6) for 1 h at room temperature and incubated with primary antibodies overnight at 4°C against either Myc, GFP, eIF5A, hyp-eIF5A, HA, H4, PGK1 or glyceraldehyde-6-phosphate dehydrogenase. Detailed information of the antibodies used in this study can be found in Table S4. Bound antibodies were detected using the appropriate horseradish peroxidase-conjugated secondary antibodies. Chemiluminiscent signals were detected with an ECL Prime Western blotting detection kit (GE Healthcare) and digitally analyzed using ImageQuant LAS 4000 software (GE Healthcare). In order to capture variation across all samples, the signal in each lane was normalized to the mean signal across all lanes in a single blot. Then, the resulting signal of bands was normalized against the corresponding G6PDH resulting signal. At least three biological replicates of each sample were analyzed.

### Fluorescence microscopy and analysis

Yeast cells were grown to a logarithmic phase in YPD medium, centrifuged, washed and subjected to standard fluorescence and phase contrast microscopy. Fluorescence images were acquired using an Axio Imager Z1 fluorescence motorized microscope equipped with a Plan Apochratic x63/1.4 oil-immersion objective and a 100 W mercury lamp (Carl Zeiss, Germany). Images were recorded with an AxioCam MRm digital camera (Carl Zeiss, Germany). To study nuclei localization, cells were incubated with 1 µg/mL 4′,6-Diamidino-2-phenylindole dihydrochloride (DAPI, Thermo Fisher Scientific) for 5 mins in the dark, washed and subjected to microscope. The following excitation and emission wavelengths were used: DAPI (excitation 359 nm; emission 457 nm) and GFP (excitation 475 nm; emission 509 nm). The same exposure times were used to acquire all images and at least three biological replicates of each sample were analyzed.

All the imaging analysis was performed on Image J software. For the analysis of fluorescence intensity signal ROIs were manually outlined around cells using the "Freehand" selection tool in ImageJ on DIC images. After background subtraction, the amount of GFP-eIF5A, its different versions and eIF4e-GFP (total fluorescence intensity) was quantified in the whole cell and in the nuclei section using the corresponding DAPI staining images. At least 100 single cells were scored from three independent experiments.

**Chromatin immunoprecipitation**

For eIF5A and Rpb1 chromatin binding experiments, yeast cell cultures were grown at 25°C in YPD and transferred to 37°C for 4 h when using temperature-sensitive strains. For the cross-linking reaction, formaldehyde was added (1% final concentration) to a volume of 45 mL of culture for 15 min at room temperature with occasional inversion. Then, the reaction was stopped with glycine (0.14 M) incubation for 5 min at room temperature. The chromatin immunoprecipitation (ChIP) experiments were performed as previously described (Li et al., 2016) with the following modifications: after reversing formaldehyde-mediated cross-linking, samples were treated with proteinase K, and DNA was purified using the GeneJET PCR Purification Kit (Thermo Fisher Scientific, #K0702) according to the manufacturer's instructions. To determine the enrichment of the DNA regions bound by the protein of interest, qPCR was run as described above using the primers listed in Table S3. The qPCR amplification data were normalized with the total input DNA value in the corresponding whole cell extract. Detailed information of the antibodies and dynabeads used can be found in Table S4.

At least three biological replicates of each sample were analyzed.

**ChIP-seq and sequencing analysis**

Wild-type yeast cells exponentially grown in YPD at 25°C were used for ChIP-seq experiments. The concentration of the DNA samples (inputs and IPs) was quantified with Qubit dsDNA HS kit (Invitrogen) and fragment size distribution of the inputs was assessed on a TapeStation using the D5000 HS assay (Agilent). Libraries for ChIP-Seq were prepared at IRB Barcelona Functional Genomics Core Facility. Briefly, dual-indexed DNA libraries were generated from 2.1 – 3.1 ng of DNA samples using the NEBNext Ultra II DNA Library Prep kit for Illumina (New England Biolabs). 16 cycles of PCR amplification were applied to all libraries.

The final libraries were quantified using the Qubit dsDNA HS assay and quality controlled with the Bioanalyzer 2100 DNA HS assay (Agilent). An equimolar pool was prepared with the four libraries and submitted for single end 50 nt sequencing on a NextSeq2000 (Illumina). More than 4 Gbp of reads were produced, with a minimum of 17 million of single end reads per sample.

Data analyses was performed at the Statistical and Omics Data Analyses facility of the SCSIE-Universitat de València. Raw reads were quality-checked and trimmed to remove adapters and low-quality bases (Phred score ≥ 28 and reads ≥ 40 bp) using fastp (v0.23.1) (1) and FastQC (v0.11.5) (2) tools. Processed reads were aligned using Hisat2 (version v2.2.1) (3) and S. cerevisiae genome (R64-1-1) (Ensembl release 110). Alignments were processed by SAMtools (version v1.13) (4) and alignment quality will be evaluated with QualiMap (v2.2.2d) (5). HTSeq v2.0.4 was used to generate raw counts per gene, using the 'gene_id' attribute from exon features in the GTF annotation file (Saccharomyces_cerevisiae.R64-1-1.110.gtf) (6). Enrichment graphic were obtained with ngs.plot (version v2.61) (7).

The eF5A binding value for each individual gene was calculated as the relative value respect the mean value for all genes.

Biological process Gene Ontology (GO) terms overrepresented in genes with the highest eIF5A binding or the highest eIF5A translation dependence measured with the Protein Pause Index (PPI). Gene set enrichment analysis was done using the ReviGO software.

**Protein Pause Index (PPI) calculation**

The dependency of each yeast gene on eIF5A for translation of its mRNA was estimated by calculating the PPI. First, the number of the top 43 eIF5A-dependent tripeptide motifs present in the encoded protein amino acid sequence was determined. These motifs cause ribosome pausing when eIF5A is depleted in yeast cells, as described by Pelechano and Alepuz (2017). Secondly, the number of each motif was multiplied by its pause strength value, as revealed by 5PSeq analysis (Pelechano and Alepuz, 2017). Thirdly, the PPI for each gene was obtained as the sum of motifs x strength.

**Determination of individual transcription rates and mRNA levels**

To determine the synthesis rate (TR) in the corresponding strains, a genomic run-on (GRO) was performed as originally described in (García-Martínez et al., 2004). Briefly, wild-type and *tif51A-1* cells were grown to early mid-log phase at 25°C and then transferred to 37°C for 4 h. Culture volumes were adjusted so that approximately $6x10^8$ cells were harvested to perform the run-on. A second sample corresponding to 20 mL of cell culture was also harvested for total RNA extraction. Both pellets were flash frozen and stored at -20°C. Upon defrosting, the cell pellet was washed in cold water and resuspended in 1 mL of 0.5% cold N-lauryl sarcosine sodium sulfate (sarkosyl) and transferred to a new tube. After permeabilization, cells were centrifuged at 6000 rpm for 1 min and the supernatant was removed. To perform the run-on, cells were resuspended with 115 µL of distilled water, 120 µL of 2.5x transcription buffer, 16 µL of ACG mix, 6 µL of 0.1 M DTT and 20 µL of [α-33P]-UTP (3000 Ci/mmol). The mixture was incubated at 30°C for 5 min and 600 rpm shaking to allow transcription elongation. The transcription was stopped with cold distilled water and the cells were collected by centrifugation at 6000 rpm for 1 min to remove the non-incorporated radioactive nucleotide. Then, total RNA was isolated following the phenol:chloroform protocol. A 5 µL aliquot was used for specific radioactivity determination using the Tricarb scintillation counter (Perkin Elmer). Home-made macroarray nylon filters (García-Martínez et al., 2004) were pre-hybridized for 1 h with hybridization solution at 65°C. 300 µl of 2x hybridization solution were added to all the *in vivo* labelled RNA samples and mixed with 3 mL of hybridization solution. Macroarray filters were hybridized for 48 h in a roller oven at 65°C. The macroarrays were vacuum-sealed with plastic and exposed to an imaging plate (BAS-MP, FujiFilm) for the desired time depending on the signal intensity (1-7 days). The imaging plate was read at 50 µm resolution in a phosphorimager scanner (FLA-3000, FujiFilm).

For the second cell aliquot, total RNA was isolated following the phenol:chloroform protocol. Approximately 50 µg of RNA were purified using the Quiaquick kit (Qiagen) following manufacturer's instructions and used for reverse transcription into cDNA. For that, 200 U of Maxima Reverse Transcriptase (200 U/µL), 3 µL of oligo d(T)15VN (500 ng/µL), 1 µL of RNaseOUT, 3 µL of 0.1 M DTT, 6 µL of 5x RT buffer, 1.5 µL of dNTP mix, and 4 µL of [α-33P]-dCTP (3000 Ci/mmol) were added to a final volume of 30 µL. The labelling reaction was incubated for 2 h at 50°C and the reaction was stopped by adding 1 µL of 0.5 M EDTA. The labelled cDNAs were purified by a S300-HR MicroSpin column (Amersham Biosciences) so the non-incorporated radioactive nucleotide was removed. The hybridizations were performed as described previously for GRO except that cDNA samples were denatured at 95°C for 5 min prior hybridization and a final concentration of 3.5 x 106/mL was employed. Macroarray filters were hybridized for 24 h in a roller oven at 65°C. Both GRO and cDNA labelled samples belonging to the same sampling were successively hybridized against the same filter.

For the quantification of hybridization signals and subsequent analysis procedures: the experiments were always done in triplicate to reduce the variability provided by differences in *in vivo* incorporation, RNA extraction and hybridization. The scanned microarray images from both GRO and cDNA experiments were quantified using Array Vision software (Imaging Research) taking the sARM density (after background subtraction) as signal. Analysis of the data was performed as described (García-Martínez et al., 2004). For GRO and cDNA analysis, values that were at least 1.2 times higher than the local background were taken as valid measurements. An average data set of three replicates for each sample was created using median absolute deviation normalization by ArrayStat software (Imaging Research Inc.). Both GRO and cDNA hybridizations were normalized within each experiment replicate by the global mean procedure. Average cDNA values for each gene were finally corrected by percentage of guanines present in each probe-coding strand while average TR values for each gene were corrected by percentage of uridines present in each probe-coding strand.

The median values of the cell volumes of the population were obtained by a Coulter-Counter Z series device (Beckman Coulter, USA). We obtained the growth rates by growing 50 ml of yeast cultures in 250-ml flasks with shaking (190 rpm) at the corresponding temperature. Aliquots were taken every hour in the exponential phase and their $OD_{600}$ (from 0.05 to 1.5) were measured. The generation time in the exponential phase were estimated from growth curves.

**Chromatin association assay**

The chromatin association assay experiment was performed as previously described in (Battagliaetal., 2017), with modifications. Yeast cell cultures were grown in 300 mL of YPD medium at 25°C to mid-log phase ($OD_{600}$ 0.5). Subsequent steps were performed at 4°C with precooled buffers and in the presence of a fresh protease-inhibitor mix. Cells were collected by centrifugation, washed with 1×TBS buffer and with lysis buffer (150 mM NaCl, 50 mM HEPES-KOH pH7.5, 1 mM benzamidine, 1 mM PMSF, 1 mM EDTA, 1% TritonX-100, 0.1% sodiumdeoxycholate, 0.1% SDS and one protease inhibitor cocktail).

Cell pellets were flash frozen, thawed, resuspended in 1 mL lysis buffer, and disrupted via bead beating (FastPrep−24 Instrument, MP Biomedicals, LLC., France) in the presence of 0.5 mL of glass beads (MERCK, USA) for 40s at 4m/s, followed by an incubation of the sample for 1 min on ice. This was repeated eight times. The lysate was divided into two samples. One half was treated with 15 U of RNase A and 15 U of RNase T1 (Ambion, UK); the other half was treated with the same volume of the RNase storage buffer (10 mM HEPES pH 7.5, 1 mM EDTA, 0.1% Triton X-100, 50% glycerol). After 1 hour incubation at 25ºC, chromatin was isolated by centrifugation at 13,000 rpm for 20 min. This was repeated three times.

Chromatin was solubilized in 200 µL lysis buffer via sonication with a Bioruptor Standard Water bath Sonicator instrument (Diagenode). Chromatin solutions were then analysed by SDS-PAGE and Western blotting against eIF5A, cytoplasmic PGK1 and nuclear H4 proteins with specific antibodies. Detailed information of the antibodies used can be found in Table S4. At least three biological replicates of each sample were analyzed.
